# Cytoskeleton members, MVBs and the ESCRT-III HvSNF7s are putative key players for protein sorting into protein bodies during barley endosperm development

**DOI:** 10.1101/595108

**Authors:** Valentin Roustan, Julia Hilscher, Marieluise Weidinger, Siegfried Reipert, Azita Shabrangy, Claudia Gebert, Bianca Dietrich, Georgi Dermendjiev, Pierre-Jean Roustan, Eva Stoger, Verena Ibl

## Abstract

Cereal endosperm is a short-lived tissue adapted for nutrient storage, containing specialized organelles, such as protein bodies (PBs) and protein storage vacuoles (PSVs), for the accumulation of storage proteins. PBs can be used as efficient biotechnological systems to produce high yields of stable recombinant proteins. During development, protein trafficking and storage require an extensive reorganization of the endomembrane system. Consequently, endomembrane-modifying proteins will influence the final grain quality, yield and recombinant protein production. Barley, a cereal crop of worldwide importance for the brewing industry, animal feed and to a lesser extent, human nutrition, has been identified as promising candidate for recombinant protein production. However, little is known about the molecular mechanism underlying endomembrane system remodeling during barley grain development. By using *in vivo* label-free quantitative proteomics profiling, we quantified 1,822 proteins across developing barley grains. Based on proteome annotation and a homology search, 95 proteins associated with the endomembrane system were identified, and 83 of these exhibited significant changes in abundance during grain development. Clustering analysis allowed characterization of three different development stages; notably, integration of proteomics data with *in situ* subcellular microscopic analyses showed a high abundance of cytoskeleton proteins associated with acidified protein bodies at the early development stages. Endosomal sorting complex required for transport (ESCRT)-related proteins and their transcripts are most abundant at early and mid-development. Specifically, multivesicular bodies (MVBs), and the ESCRT-III HvSNF7 proteins are associated with protein bodies (PBs) during barley endosperm development. Taken together, our proteomics results specifically identified members of the cytoskeleton, MVBs, and ESCRT as putative key players for protein sorting into PBs during barley endosperm development. These results present a comprehensive overview of proteins involved in the rearrangement of the endomembrane system during barley early grain development and will provide the basis for future work on engineering the endomembrane system to optimize nutrient content and to produce high yields of recombinant proteins.

## Background

After differentiation, fully developed cereal endosperm can account for up to 75% of the grain weight and comprises four major cell types: aleurone, starchy endosperm, transfer cells, and the cells of the embryo surrounding region (Olsen, 2001). The starchy endosperm thereby functions as a storage site, as it accumulates starch and seed storage proteins (SSPs) (Olsen, 2004). The aleurone layer supports seed germination by mobilizing starch and SSP reserves in the starchy endosperm by releasing hydrolytic enzymes that help to degrade the stored nutrients in the endosperm (Olsen, 2004). Contrary to the persistent endosperm of cereals, the cellular endosperm of *Arabidopsis thaliana* supports the developing and growing embryo, resulting in a gradually depleted endosperm as the embryo grows. Finally, the massive *A. thaliana* embryo is only accompanied by a single peripheral layer, the aleurone layer, in mature seeds (Olsen, 2004). Consequently, *A. thaliana* cannot to be used as a model system to study the endomembrane system in grain endosperm.

In cereals, SSPs, which account for more than 50% of the grain protein content (Psota et al., 2007; Guo et al., 2016), accumulate in the outer layer of the endosperm, the subaleurone, and in the starchy endosperm, the latter in parallel with starch granules (Moore et al., 2016). In barley, for instance, globulins and prolamins comprise the major endosperm SSPs (Shewry and Halford, 2002).

The SSP trafficking routes depend on the cereal species, endosperm layer and development stage (Ibl and Stoger, 2012; Arcalis et al., 2014; Zheng and Wang, 2014). SSPs are produced by the secretory pathway and reach their final destinations by two main routes: soluble albumins and globulins travel through ER and Golgi to protein storage vacuoles (PSVs); and most prolamins are finally deposited in specific, ER-derived protein bodies (PBs) or/and in PSVs. Along with these main routes, other organelles are proposed to be involved in the SSP trafficking in cereal endosperm, e.g., multivesicular bodies (MVBs), precursor-accumulating (PAC) vesicles and/or dense vesicles (DVs) (Ibl and Stoger, 2012).

More recently, PBs have arisen as potential targets for recombinant protein production, especially pharmaceutical proteins (Tschofen et al., 2016). Indeed, these specialized storage organelles provide the ability to accumulate and store a high amount of (recombinant) proteins, subsequently offering high stability *in planta* and during harvest (Stoger et al., 2005; Hofbauer and Stoger, 2013). Interestingly, PBs are highly resistant to gastrointestinal digestion *in vitro* and *in vivo*, enabling PB bioencapsulation (reviewed in (Arcalis et al., 2014). In this context, barley *(Hordeum vulgare)*, the fourth-most important cereal in terms of food production after maize, wheat, and rice (FAOSTAT 2016), holds great molecular farming potential. Barley is a self-pollinating, diploid crop and can be easily transformed (Horvath et al., 2000; Eskelin et al., 2009; Magnusdottir et al., 2013; Ritala et al., 2014). Recently, active mammalian antimicrobial LL-37 peptide was produced in barley endosperm (Holaskova et al., 2018). Further, recombinant anti-HIV-1 monoclonal antibody 2G12 was produced and accumulated within the endosperm of transgenic barley grain; 2G12, stored in ER-derived protein bodies, showed a comparable equilibrium and kinetic rate constants compared to recombinant 2G12, synthesized in Chinese hamster ovary cells (Hensel et al., 2015).

Depending on the protein trafficking, PB composition is affected thereby influencing the qualitative output. For instance, the SSP composition defines the malting purposes, most specifically for the brewing industry (Gupta et al., 2010). Thus, endomembrane-modifying proteins within the endomembrane system will have an influence in the final grain quality/yield and recombinant protein production.

In barley, microscopic analyses revealed that spherical PSVs, which are stable in the aleurone during development, underwent a dynamic rearrangement including fusion, rupture and degeneration in the subaleurone and starchy endosperm (Cameron-Mills, 1980; Ibl et al., 2014; Hilscher et al., 2016). Between 8 and 12 DAP, PSVs reduced their size and even degenerated in the starchy endosperm (Ibl et al., 2014). Simultaneously, PBs were found in distinct compartments in the subaleurone but also within the vacuoles, where fusion events of PBs were observed by live cell imaging (Ibl et al., 2014). In the starchy endosperm, PBs were tightly enclosed by vacuoles that finally degenerated and released the PBs (Ibl et al., 2014). Recent bioinformatic, proteomic and RT-qPCR analyses shed the first light on proteins possibly involved in the rearrangement of the endomembrane system in barley endosperm (Hilscher et al., 2016; Roustan et al., 2018; Shabrangy et al., 2018): ESCRT-III proteins were identified by a bioinformatic assay (Hilscher et al., 2016) and localization studies of recombinantly expressed HvSNF7a, HvVPS24 and HvVPS60a ESCRT III components in barley revealed different localization of these proteins within the endosperm (Hilscher et al., 2016). It was further suggested that the steady-state association of ESCRT-III may be influenced by cell layer–specific protein deposition or trafficking and remodeling of the endomembrane system in the endosperm (Hilscher et al., 2016). Additionally, the ER was identified to be most abundant in the starchy endosperm and ER rearrangements were characterized, including re-localization of HvPDIL1-1 during development (Roustan et al., 2018).

This work aimed to temporally map *in situ* the endomembrane system during barley endosperm development, using integrative cell biology experimental approaches. In this study, *in vivo* label-free proteomics approaches allowed the quantification of 1,822 proteins. Among these, 95 proteins could be associated with the endomembrane system. More specifically, seven out of eight proteins related to the ESCRT machinery showed specific expression patterns and transcript abundances associated with early and mid-development. In this context, confocal and transmission electron microscopy analyses located MVBs and HvSNF7 at the periphery of PBs and later within PBs, playing a putative role in protein sorting to PBs at mid-development.

Taken together, our proteomics results provide comprehensive developmental mapping of proteins potentially involved in the rearrangement of the endomembrane system during the barley early grain-filling process and the specific identification of cytoskeleton members, MVBs, and ESCRT as putative key players for protein sorting into PBs during barley endosperm development.

## Materials and methods

### Plant material and growth conditions

Barley (*H. vulgare L.*) wild-type variety Golden Promise (GP) and transgenic lines (TIP3-GFP, p6U::SNF7.1-mEosFP) were cultivated as described in (Ibl and Stoger, 2014). Caryopses were harvested at different stages of grain development including 6–8, 10, 12-18 and ≥ 20 days after pollination (DAP) (designated as 6, 10, 12 and ≥ 20 DAP) of three biological replicates.

### Sample preparation for proteomics analyses

Total proteins were extracted from barley grains harvested at 6, 10, 12 and ≥ 20 DAP in three biological replicates. Stages were chosen as previously described (Shabrangy et al., 2018). Extraction was performed following an adapted phenol-phase extraction protocol as described in (Roustan et al., 2017). Subsequently, proteins were re-suspended in urea buffer (8 M urea, 100 mM ammonium bicarbonate) to measure protein concentration with a Bradford Assay prior protein content normalization and trypsin digestion. Following overnight digestion, peptides were desalted using a C18 solid phase extraction (Agilent Technologies, Santa Clara, USA). After solid-phase extraction, the corresponding eluates were dried in a vacuum concentrator.

### Nano-HPLC and Orbitrap Elite tune methods

Mass spectrometry (MS) was performed as previously described (Roustan et al., 2018). Peptide pellets were resolved at a protein concentration equivalent of 0.1 µg/µL. A total of 0.5 µg of the mixture was separated on an EASY-Spray PepMap RSLC 75 μm × 50 cm column (Thermo Fisher Scientific, Waltham, Massachusetts, USA). After elution using a 150 min linear gradient at a 300 nL/min flow rate generated with an UltiMate 3000 RSLCnano system, the peptides were measured with an LTQ-Orbitrap Elite (Thermo Fisher Scientific, Waltham, Massachusetts, USA) as described in (Roustan et al., 2018). Mass analyzer settings were as follows: ion transfer capillary temperature 275 °C, full scan range 350–1800 m/z, FTMS resolution 60,000. Each FTMS full scan was followed by up to twenty data-dependent (DDA) CID tandem mass spectra (MS/MS spectra) in the linear triple quadrupole (LTQ) mass analyzer. Dynamic exclusion was enabled using list size 500 m/z values with an exclusion width ±10 ppm for 60 s. Charge state screening was enabled and unassigned and +1 charged ions were excluded from MS/MS acquisitions. For injection control, automatic gain control for full scan acquisition in the Orbitrap was set to 1 x 106 ion population, and the maximum injection time (max IT) was set to 200 ms. Multistage activation was enabled with neural losses of 24.49, 32.66, 48.999, 97.97, 195.94, and 293.91 Da for the 10 most intense precursor ions. Prediction of ion injection time was enabled, and the trap was set to gather 5 x 103 ions for up to 100 ms. Orbitrap online calibration using internal lock mass calibration on m/z 371.10123 from polydimethylcyclosiloxane was used.

### Data processing and protein identification

Raw files were processed with MaxQuant 1.5 (http://www.maxquant.org) and the Andromeda search algorithm (Cox and Mann, 2008; Cox et al., 2011; Tyanova et al., 2015) on the barley UniProt database (http://uniprot.org). Peptide identification was performed using the following settings: mass tolerance for precursors was set to 5 ppm and for fragment masses up to 0.8 Da. The maximum FDR was set to 0.01%. Three missed cleavages were allowed. The dynamic modifications allowed were: methionine oxidation (M) and protein N-terminal acetylation. The fixed modification allowed was: Carbamidomethyl (C). Label-free quantification was done at the MS1 level with at least two peptides per protein. PTXQC was used to assess data quality and statistical analysis was performed with Perseus 1.5 software (Bielow et al., 2016; Tyanova, 2016).

Protein annotation was performed with the MERCATOR tool (http://mapman.gabipd.org/de) (Lohse et al., 2014). Unknown proteins were identified by using BLAST at the UniProt homepage searching for the most identical cereal protein. Proteins were classified to “compartment-specific proteins” (functional associated with a specific subcellular endomembrane pathway or organelle), and as “trafficking regulators” (functional associated with several organelles) based on published data. In the final dataset, representative proteins were quantified in at least 9 of the 12 samples analyzed. Data were first Log2 transformed prior Z-scores (zero mean, unit variance) and were finally used to calculate the relative protein abundance. A one-way analysis of variance (ANOVA) and Student’s t-tests were performed with Perseus 1.5 software. Cluster analysis was performed with fuzzy-c means algorithm implemented in GProX (Rigbolt et al., 2011). Generation of protein-protein interaction (PPI) networks was conducted via the Search Tool for the Retrieval of Interacting Genes/Proteins (STRING) database for known and predicted protein-protein interactions (http://string-db.org/) with default parameters (Franceschini et al., 2013). The MS proteomic data have been deposited to the ProteomeXchange Consortium (Deutsch et al., 2017) via the PRIDE (Vizcano et al., 2016) partner repository with the dataset identifier PXD009722.

### Western blots

To confirm the MS results, we have used the same extraction protocol for the Western blots as for the MS analysis. For the analyses of the transgenic line p6U::SNF7.1-mEosFP, five transgenic and wild type (GP) grains were extracted by PBS buffer pH 6. After protein extraction, we measured the quantity of protein with a Bradford assay. Western blots were performed as previously described (Roustan et al., 2018). A total of 2 µg of PageRuler Plus Prestained Protein Ladder (#26 619, Thermo Fisher Scientific, Waltham, Massachusetts, USA) and 20 µg of the samples were loaded on a 10% acrylamide gel. Antibodies were diluted as following: HSP70 Heat shock protein (#AS 08 371, Agrisera, Vännäs, Sweden), 1/3000; polyclonal rabbit anti-SNF7 antibody (kindly provided by (Teis et al., 2010), 1/1000; and Amersham ECL Rabbit IgG, HRP-linked whole Ab (from donkey) (#NA934VS, GE Healthcare, Illinois, Chigaco, United States), 1/10,000. The ECL prime Western blotting detection reagent (#RPN2232, GE Healthcare, Illinois, Chigaco, United States) was used for development.

### RT-qPCR Analysis

RT-qPCR analysis of *ESCRTs* was performed according to the MIQE guidelines (Bustin et al., 2009). RNA was isolated from grains from four development stages: 6, 10, 12 and ≥ 20 DAP. The RNA concentration was measured at 260 nm using a UV spectrophotometer (NanoDrop Technologies, Thermo Fisher Scientific, Massachusetts, USA) and ranged from 70 to 224 ng/μL. RNA integrity was assessed by microfluidic capillary gel electrophoresis using the ExperionTM RNA HighSens Analysis Kit (#7007105, Bio-Rad Laboratories, Hercules, Carlifornia, USA) with Experion software 3.2P. The quality of RNA index (QRI) showed acceptable quality of the isolated RNA. Next, cDNA was synthesized as recently described in (Shabrangy et al., 2018). Specific primers were designed for HvPDIL1-1 using Primer-Blast (https://www.ncbi.nlm.nih.gov/tools/primer-blast/) (see Supplemental Table 1).

Besides bioinformatic analyses, the specificity of the primers was tested by PCR on the cDNAs. At least three biological replicates were used, and three technical replicates were performed for RT-qPCR. For the normalization studies of *ESCRT* transcripts, we used the following reference genes as described in Shabrangy et al. (2018): *ARF* (ADP-ribosylation factor), *FBPA* (fructose-bisphosphate aldolase), and *SAM* (S-adenosyl-L-methionine) for whole grains. Normalization was calculated as described in (Vandesompele et al., 2002; Shabrangy et al., 2018). For statistical analyses we performed a Student’s t-test (two-tailed distribution, two-sample unequal variance (heteroscedastic)) using the Microsoft Excel software program.

### Cloning of constructs

#### Bimolecular Fluorescence Complementation (BiFC)

The backbone of all vectors (MKK4_SPYCE and MPK3_SPYNE, kindly provided by Dr. Andrea Pitzschke) used in this study contain a p35S promoter (Cauliflower Mosaic virus 35S promoter), 5’UTR (untranslated region from tobacco etch virus), our genes of interest (Hvsnf7.1), C-terminal or N-terminal sequence of YFP (SPYCE or SPYNE, t35 (Cauliflower Mosaic virus 35S terminator), HA-tag (Human influenza hemagglutinin) or c-MYC-tag, and a kanamycin antibiotic resistance sequence. *HvSNF7.1* (according to *HvSNF7a.1* described in (Hilscher et al., 2016) was cloned into the vector pCR2.1 (#K200001, Thermo Fisher Scientific, Massachusetts, USA) using HvSNF7.1_NcoI-F and HvSNF7.1_NotI-R as primers, digested by NcoI and NotI and inserted into previously digested MKK4_SPYCE and MPK3_SPYNE, respectively. To obtain a pSPYCE vector without insert for control reactions, plasmids were cut (NcoI/NotI), blunted using Klenow fragment and re-ligated. All the clones were verified by sequencing.

#### Yeast two-hybrid (Y2H)

The restriction sites NdeI and Sfil were introduced into HvSNF7.1 by PCR using the primers pGADT7_SNF7F and pGADT7_SNF7R3. Using NdeI and Sfil, HvSNF7.1 was ligated into the vector pGEM®-T Easy (#A1360, Promega, Madison, Wisconsin, United States). All the clones were verified by sequencing and finally cloned into the target vectors pGADT7 AD (#630442, Clontech Laboratories, Mountain View, California, United States) and pGBKT7 DNA-BD (#630443, Clontech Laboratories, Mountain View, California, United States). Positive clones were verified by sequencing.

#### p6U::SNF7.1-mEosFP

To obtain HvSNF7-mEosFP driven by the barley hordein D promoter (p6U) and ended by the nopaline synthase terminator, HvSNF7.1-mEosFP (previously described in (Hilscher et al., 2016) was ligated in the following into the vector p6U_pHordeinD (kindly provided by Eszter Kapusi, unpublished) using MluI. The p6U vector backbone originates from DNA Cloning Service e.K. (http://www.dna-cloning.com/vectors/Binaries_hpt_plant_selection_marker/p6U.gbk). The sequence of the Hordein D promoter was taken from (Horvath et al., 2000). The vector p6U_pHordeinD is based on p6U_pHordein_BamHI_SP), which was cut using BamHI/HindIII, blunted using Klenow fragment and re-ligated.

#### YFP-FYVE

The plasmid 2xCaMV35S::YFP-2xFYVE harboring the YFP-FYVE marker was a gift of Teun Munnik (Vermeer et al., 2006).

### Transformation of barley endosperm cells

Barley (GP) transformation was carried out using particle bombardment (Wan and Lemaux, 1994; Ibl and Stoger, 2014). T1 plants surviving hygromycin selection were genotyped using primer pairs as described in Supplemental Table1. Homozygous T4 grains were used from transgenic p6U::SNF7.1-mEosFP plants for microscopic and Western blot analyses. The intact fusion protein SNF7.1-mEosFP was detected by Western blot using polyclonal rabbit anti-SNF7 antibody (kindly provided by D. Teis), which could detect the transgenic SNF7.1-mEosFP (49 kDa), wild type SNF7.1 (25 kDa), and SNF7.2 (31 kDa) (Supplemental Figure 1).

### Microscopy

#### Live cell imaging

GP and the transgenic line TIP3-GFP (Ibl et al., 2014) were used for live cell imaging as previously described (Ibl et al., 2014). ER-Tracker™ Green (BODIPY™ FL Glibenclamide; #E34251, Thermo Fisher Scientific, Waltham, Massachusetts, USA) and LysoTracker™ Red DND-99 (#L7528, Thermo Fisher Scientific, Waltham, Massachusetts, USA) were used to visualize ER and acidic compartments, respectively. ER-Tracker™ Green was used as previously described (Ibl et al., 2014). FM™ 4-64 Dye (N-(3-triethylammoniumpropyl)-4-(6-(4-(diethylamino)phenyl) hexatrienyl) pyridinium dibromide, #T13320, Thermo Fisher Scientific, Waltham, Massachusetts, USA) was used as an endocytic tracer (Bolte et al., 2004). In detail, at least three randomly selected transgenic and GP grains were harvested at 6, 12 and ≥ 20 DAP, sectioned, washed, and stained as follows: FM4-64, 10 minutes (final concentration, 8 µM, diluted from 1 mM DMSO stock with water); ER-Tracker™ Green, 1 h (final concentration, 2 µM, from 1 mM DMSO stock with water); LysoTracker™ Red, 30 min (final concentration, 2 µM, from 1 mM DMSO stock with water). Mock treatments included the final DMSO concentration in water. Sections were mounted in tap water and immediately imaged by the Leica SP5 CLSM using sequential scans with filter settings for GFP (excitation 488 nm, emission 500–530 nm), LysoTracker™ Red (excitation 561 nm, emission 570–630 nm), and ER-Tracker™ Green (excitation 488 nm, emission 500–531 nm). Transgenic p6U::SNF7.1-mEosFP grains were chipped, mounted in tap water and analyzed by CLSM with excitation at 488 nm, emission 508–540 nm.

#### Bimolecular Fluorescence Complementation (BiFC)

A single colony of transformed *Agrobacterium tumefaciens* was inoculated in 5 mL of YEB-Medium (0.5% beef extract, 0.5% sucrose, 0.1% yeast extract, 0.05% MgSO_4_*7H_2_O) containing appropriate antibiotics and incubated at 28 °C overnight. In the morning, 1 ml of the pre-culture was taken and re-inoculated into 5 ml of the same medium. Cells were collected by centrifugation at 5000 *g* for 5 min and the pellet was washed with 1 mL infiltration buffer (10 mM MES pH 5.7, 10 mM MgCl_2_, 100 µM acetosyringone). Washed cells were re-collected by centrifugation (5000 *g*, 5 min) and washed two additional times with 500 µl infiltration buffer and finally adjusted to an OD_600_ of 0.3. The resuspended bacteria containing the corresponding binary expression vectors for BiFC were mixed in a ratio of 1:1 and incubated for 3 h in darkness. *Nicotiana benthamiana* plants were cultivated in the greenhouse on soil, maintained at 60% humidity, with a 14 h light period and a 25 °C day/19 °C night temperature cycle. The bacterial solution was inoculated into the entire leaf area through the abaxial sides using a 1 mL syringe; two leaves per plant were infiltrated. After infiltration, the plants were kept in a tray with a hood at 25 °C. After 2–5 days, the detection of protein–protein interaction by BiFC was performed using confocal microscopy (Leica SP5 CLSM). The excitation wavelength was 514 nm (argon laser) and emission was detected between 525–600 nm and 680–760 nm for YFP and autofluorescence detection, respectively.

#### Histological and immunofluorescence studies

At least three randomly selected GP grains were harvested at 6, 12 and ≥ 20 DAP and fixed, embedded and sectioned as described in (Roustan et al., 2018; Shabrangy et al., 2018). The 1.5 µm sections on glass slides were stained with toluidine blue (0.1%) for 30 s at 80 °C on a hot plate and rinsed with distilled water. Immunofluorescence microscopy of developing barley grains was performed as described by (Shabrangy et al., 2018) using: polyclonal rabbit anti-V-ATPase antibody (#AS 07 213, Agrisera, Vännäs, Sweden), dilution 1:100; polyclonal rabbit anti-actin antibody (#AS 13 2640, Agrisera, Vännäs, Sweden), dilution 1:50; polyclonal rabbit anti-tubulin-α antibody (dilution 1:50); and polyclonal rabbit anti-SNF7 antibody (dilution 1:100, kindly provided by D. Teis). Goat Anti-Rabbit IgG (H+L) Cross-Adsorbed Secondary Antibody, Alexa Fluor 488 (#A-11008, Thermo Fisher Scientific, Waltham, Massachusetts, USA) (dilution 1:30) was used as a secondary antibody. At least three sections were analyzed, and pictures captured by Nikon Eclipse Ni. Images were processed using Leica confocal software version 2.63, ImageJ and Adobe Photoshop CS5. Red channels (FM4-64) were visualized in magenta.

#### Transmission electron microscopy (TEM)

The coat was removed for grains harvested at ≥ 20 DAP, chopped and immediately fixed in 4% (w/v) paraformaldehyde plus 2.5% (v/v) glutaraldehyde in 0.1 M sodium cacodylate, pH 7.4, overnight at 4 °C. After washing with sodium cacodylate buffer, the samples were immersed in OsO_4_ for one and a half hours at room temperature followed by washing with buffer. The chemically fixed samples underwent dehydration in an ethanol series (30%, 50%,70%, 95% ethanol for 10 min each, and two times 100% ethanol). Prior to infiltration with resin, the ethanol was replaced by 100% acetone. The dehydrated samples were infiltrated with low viscosity resin (Agar Scientific Ltd, Stansted, UK) as follows: 1/3 volume resin and 2/3 volume acetone for 15 min, 1/2 volume resin and 1/2 volume acetone for 30 min, 2/3 volume resin and 1/3 volume acetone for 3 h, and pure resin overnight. Subsequently, samples were transferred in resin-filled tubes and polymerized in an oven at 65 °C for two days.

Ultrathin sections (90 nm) of the grains were cut by using an ultramicrotome LEICA EM UC7 (Wetzlar, Germany) and a diamond knife type “ultra 45°” (DIATOME Ltd., Switzerland), collected on copper grids (300 mesh) and stained with 2.5% gadolinium triacetate for 30 min and 3% lead citrate for 8 min. Grids were analyzed at 120 kV with a TEM Zeiss Libra 120 equipped with a LaB_6_ (Lanthanum-Hexaboride) cathode. Images were acquired using a side port camera Morada G2, 11 MP (Soft Imaging System GmbH, Münster, Germany) and a bottom mount camera Sharp:eye TRS (2 × 2 KP) with iTEM software and processed using Adobe Photoshop CS5.

### Yeast two-hybrid (Y2H)

The vector pGBKT7-HvSNF7.1 was co-transformed with pGADT7-HvSNF7.1 into the yeast strain Y190. In addition, pGBKT7-HvSNF7.1 was co-transformed with pGBKT7, pGBKT7 with pGADT7-HvSNF7.1 and pGBKT7 with pGADT7. Transformation was performed as following: a 3mL; preculture (YPD media, yeast extract 5 g, tryptone 10 g, glucose 10 g) was prepared from a single colony of Y190 and incubated overnight at 30 °C. About 10–15 mL of YPD media were inoculated with 1 ml preculture and incubated at 30 °C for 3 h. Cells were accumulated by centrifugation at 3,000 rpm at 20 °C for 5 min. Cells were washed with 1 mL TE buffer (10 mM Tris/HCL pH 7.5, 1mM EDTA pH 8.0) and re-centrifuged at 3,000 rpm at 20 °C for 5 min. LA-buffer (Li-Acetat*2H_2_O 1.02 g, in 100 ml TE-buffer pH 7.5) was mixed with the yeast cells and 60 µg of DNA was added. Transformation was performed adding 300 µL 50% PEG4000 (#0156.3, Carl Roth, Karlsruhe, Germany), incubated at 22 °C for 1 h, then adding 40 µL 100% DMSO and incubating at 42 °C for 10 min. Cells were collected by centrifugation and washed with TE buffer. Transformants were selected after 2 d on synthetic dropout (SD) medium lacking Leu and Trp (SD-LW) at 30 °C. To examine Y2H interactions, the transformants were streaked out on solid medium lacking Leu and Trp (SC-LW) or lacking Leu, Trp, and His (-LWH) with 5 mM 3-amino-1,2,4-triazole for 2 days at 30 °C.

### Electrolyte leakage (EL) assays

EL assays were performed as described previously (Shi et al., 2012; Jiang et al., 2017). Five grains from each development stage (6, 10, 12 and ≥ 20 DAP) were collected in 15 mL tubes containing 8 mL of deionized water, and the electrical conductivity (EC) was measured (S0). The samples were gently shaken at 22 °C for 15 min, and the resulting EC was measured again (S1). Then, the samples were incubated at 100 °C for at least 15 min and shaken at 22 °C for another 20 min, and the resulting EC was measured as S2. EL was calculated as follows:

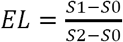

## Results

### Multivariate statistics unravel proteome reorganization across multiple biological processes during barley grain development

To identify proteins that are localized at the endomembrane system and/or are functionally associated with the rearrangement of the endomembrane system in barley, grains of different development stages including 6, 10, 12 and ≥ 20 DAP were harvested as previously described (Shabrangy et al., 2018). We used liquid chromatography-mass spectrometry (LC-MS/MS) to identify a total of 3,005 proteins. To validate our *in vivo* label-free quantification data obtained by MS, we compared the relative intensity of the heat shock protein 70 (HSP70) to a semi-quantitative Western blot using an anti-HSP70-specific antibody. Supplemental Figure 2A shows that the expression pattern of HSP70 over the four development stages is in accordance with protein quantification via MS-based proteomics (Supplemental Table 2). Thus, our biochemical data confirmed the results of our MS-based study and previously published results (Kaspar-Schoenefeld et al., 2016). Additionally, measured protein abundances were highly reproducible with an average Pearson’s correlation coefficients of > 0.95 between biological replicates (Supplemental Figure 2B). The performed Principal Component Analysis (PCA) supports these results and shows that samples grouped and clustered in a stage-specific manner (Figure 1A).

**Figure 1.**
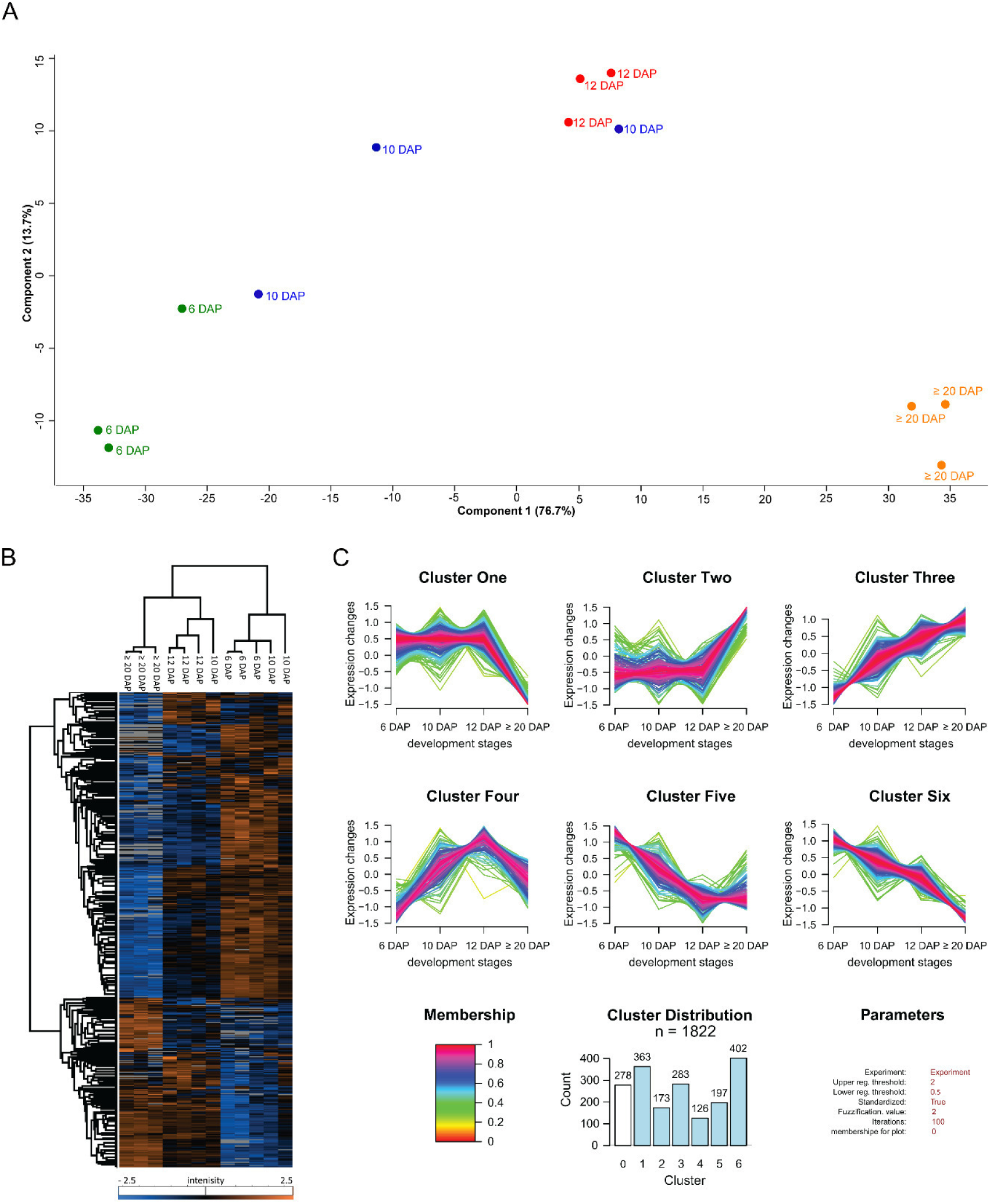
Proteome profiling during barley grain development. (**A**) PCA was conducted on logarithmically transformed protein intensities; each dot corresponds to a single biological replicate (n = 3). (**B**) Hierarchical cluster analysis of quantified proteins along barley grain development was performed with Perseus after Z-score transformation of the data (Tyanova et al., 2016). Clustering of proteins was done based on Euclidian distance while samples’ clustering is based on Pearson correlation. (**C**) Cluster of proteins dynamics along the grain development. Quantified proteins were subjected to unsupervised clustering with the fuzzy c-means algorithm implemented in GproX (Rigbolt et al., 2011). Cluster distribution indicates the number of proteins in each cluster. Membership value represents how well the protein profile fits the average cluster profile.

A total of 1,822 out of 3,005 were quantified in at least 9 out of the 12 analyzed biological samples (Supplemental Table 2). To determine the proteins that were significantly changed along the grain development, we applied a one-way ANOVA analysis, corrected with a permutation-based false discovery rate (FDR) (*p <* 0.05). Among the 1,822 quantified proteins, the abundance of 1,544 proteins was significantly changed (Supplemental Table 2), indicating that most proteins were distinctly regulated in their expression during early grain developmental stages. To assess the proteome dynamics during barley grain development, both PCA and a hierarchical bi-clustering analysis (HCA) were performed. Thereby identical clusters of stage-specific groups related to development stages of the barley grain were identified (Figure 1A and 1B). An investigation of the PCA results shows that PC1 (76.7% of variance) separated the sample based on the different development stages, particularly between the early (6 DAP) and late development stages (≥ 20 DAP). PC2 (13.7% of variance) was rather more involved in the early development stage discrimination. Protein loadings on each principal component are indicated in Supplemental Table 2. As expected, SSPs were among the highest loadings on PC1. Interestingly, most of the detected changes occurred at ≥ 20 DAP (Figure 1B).

To refine our protein expression pattern analysis, unsupervised clustering was performed with GproX software to partition the temporal profiles of 1,544 significantly changed proteins measured at all time points. Six clusters based on a fuzzy-mean clustering process (Rigbolt et al., 2011) were detected (Figure 1C and Supplemental Table 2):

Proteins presenting a higher expression level at 6 DAP than at ≥ 20 DAP belong to Clusters One, Five and Six. Those three clusters account for most of the significantly changed proteins (in total, 962 proteins). It is known that endosperm development involves cell division, cellular differentiation events and the deposition of SSPs between early, late and mid-development (Sabelli and Larkins, 2009; Zhang et al., 2016). These findings were confirmed by toluidine-stained sections prepared at 6, 12 and ≥ 20 DAP that revealed fully cellularized endosperm including three aleurone cell layers at 6 DAP (Supplemental Figure 3A). Cluster Four accounts for 126 proteins and shows its highest protein abundance between 10 and 12 DAP, which corresponds to mid-stage, where the differentiation of aleurone, subaleurone, and starchy endosperm is finalized (Figure 1C and Supplemental Figure 3A). Finally, the 456 proteins associated with Cluster Two and Three exhibit higher expression levels at ≥ 20 than at 6 DAP, corresponding to the end of mid-stage and beginning of the late stage, where the accumulation of storage reserves can be observed in the endosperm (Figure 1C and Supplemental Figure 3). Using LC-MS/MS, all 20 identified SSPs showed significantly higher abundance at ≥ 20 DAP (Supplemental Figure 3B, C).

Toluidine staining of sections prepared at 6, 12 and ≥ 20 DAP confirmed protein accumulation starting at 12 DAP and increasing at ≥ 20 DAP in subaleurone as well as in the starchy endosperm (Supplemental Figure 3A). Additionally, Cluster Two and Three contain HvPDIL1-1 and HINs (Roustan et al., 2018; Shabrangy et al., 2018). From the development stage perspective, specific trends associated with biological processes could be observed. For instance, most of the proteins related to DNA, RNA, protein synthesis, photosynthesis, and starch degradation metabolisms are highly represented in Clusters One, Five and Six. Contrary, proteins related to SSPs, Late-Embryogenesis-Associated (LEA) proteins, starch synthesis, and fermentation were preferentially associated with Clusters Two and Three (Supplemental Table 2).

### Identification of molecular regulators for the endomembrane system during barley grain development

We examined our proteomics dataset for proteins that are functionally associated with a specific subcellular endomembrane compartment (designated as “compartment-specific proteins”) as well as for spatio-regulated trafficking regulators associated with several organelles within the plant cells (designated as “trafficking regulators”). Based on a BLAST search, 95 proteins could be identified in total, where 60 compartment-specific proteins (secretory pathway, peroxisome, plasma membrane (PM), sorting, transport, degradation, and vacuolar processing) and 35 trafficking regulators were defined (dynamins, SNAREs, disulfide-generating enzyme and- carrier, ATPase, GTPase, GTPase-activating protein, RAB regulator, and GTP binding protein) within the plant cell (Table 1). The abundance of all identified endomembrane related proteins at 6, 10, 12, and ≥ 20 DAP were visualized by a heat map categorizing the protein expression pattern based on Pearson correlation (Figure 2). Compartment-specific proteins and trafficking regulators were categorized in pink and blue, respectively (Figure 2).

**Table 1.**
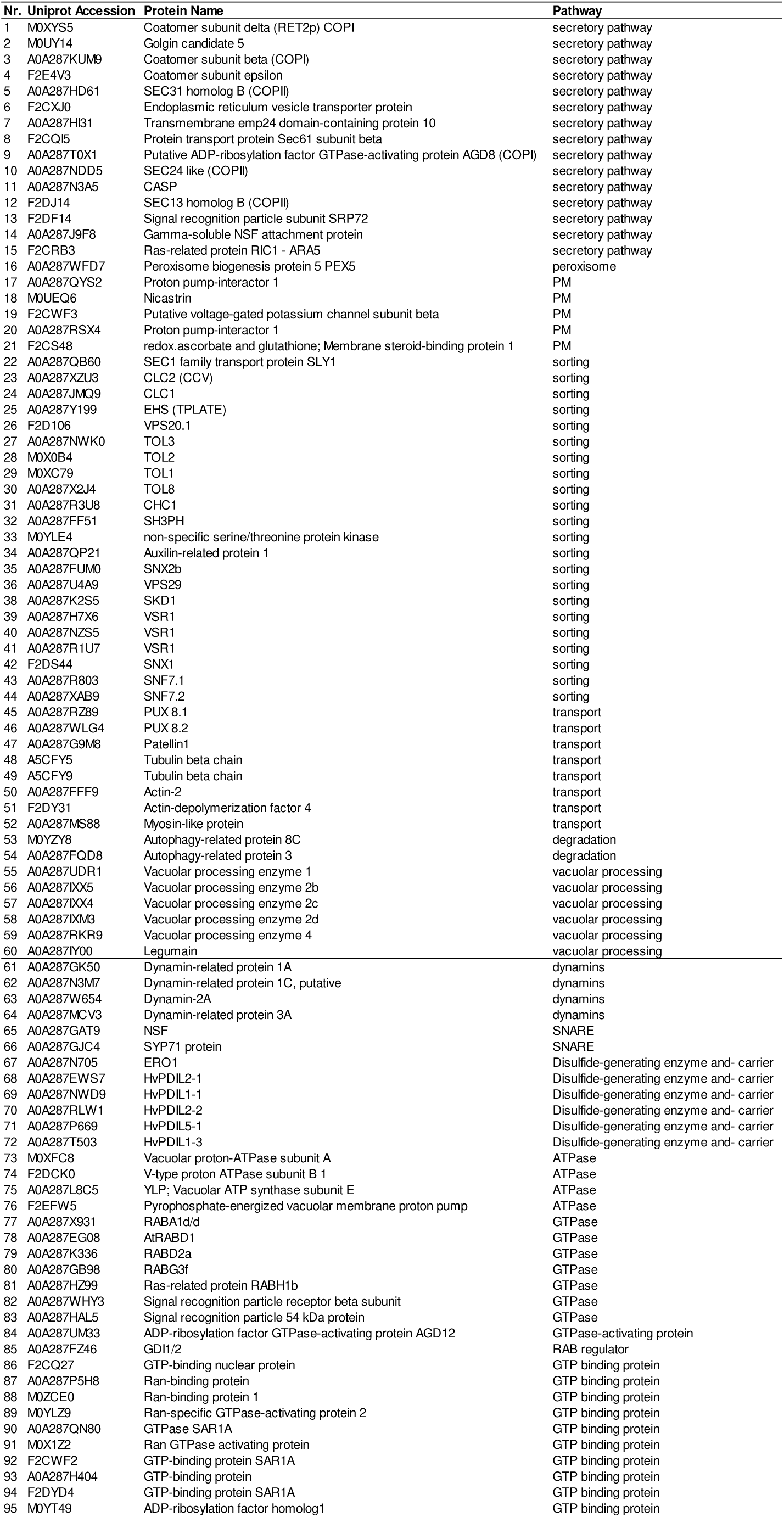
Identified proteins classified corresponding to their involvement within the endomembrane pathway and to their diverse endomembrane functions.

**Figure 2.**
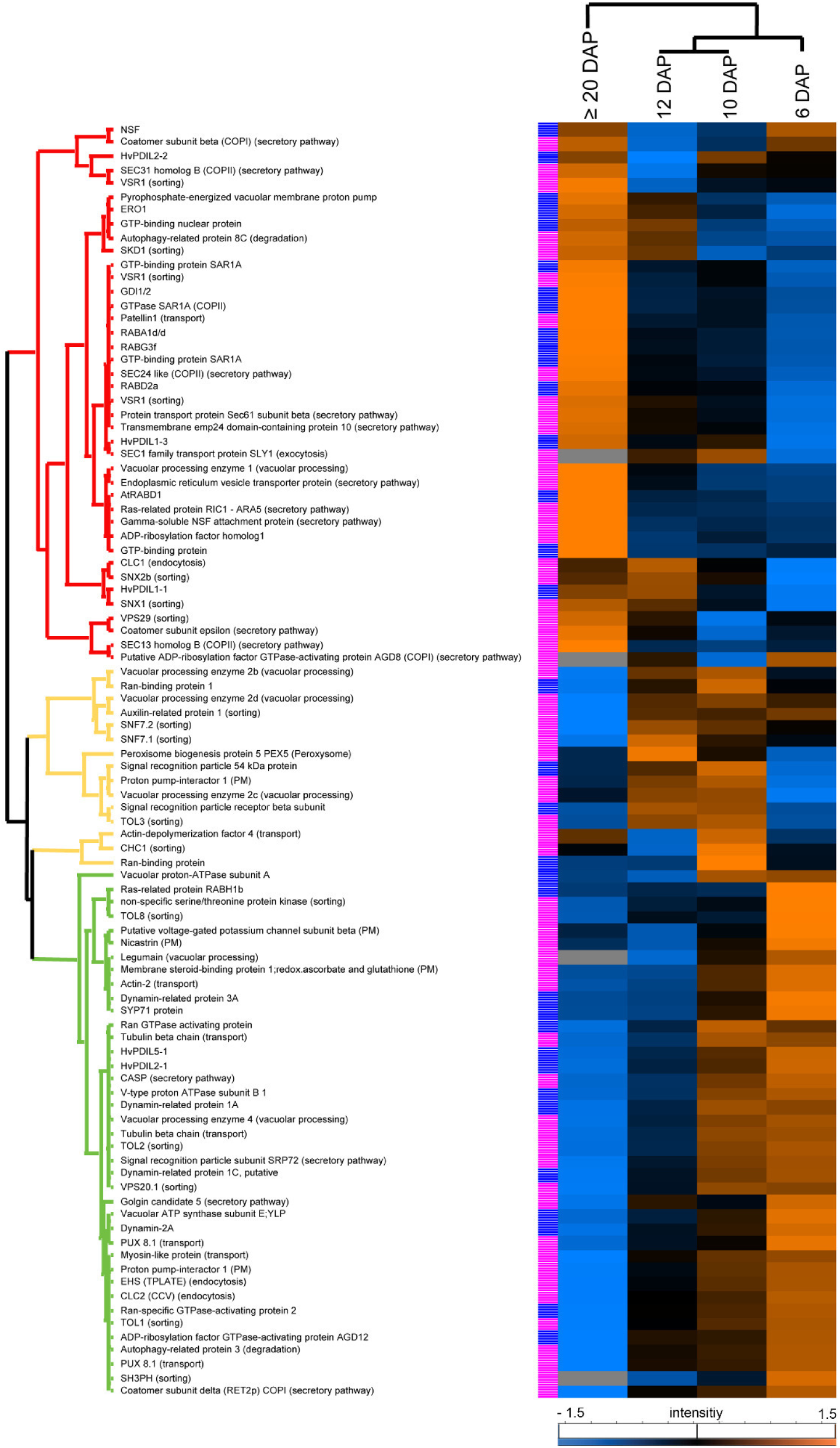
Heat map visualizes the protein expression pattern based on Pearson correlation at 6, 10, 12 and ≥ 20 DAP and clusters the proteins into three main groups: 39 proteins present a higher expression during the development stage I (green); 15 proteins presented an expression peak at development stage II (yellow); 40 proteins, associated with development stage III (red). Compartment-specific proteins and trafficking regulators are indicated with pink and blue, respectively.

Proteins could be associated with the three main development stages (Figure 2, Supplemental Figure 4A): 39 proteins present a higher expression during the development stage I (green cluster), 15 proteins presented an expression peak at the development stage II (yellow cluster), and 40 proteins are associated to the development stage III (red cluster). Within each stage, compartment-specific proteins and trafficking regulators were categorized (Table 1, Supplemental Figure 4A, B). Taken together, our data identified molecular regulators for the endomembrane system in developing barley grain.

### Development stage I contains proteins of high abundance associated with endocytosis and cytoskeleton, plasma membrane proteins, and ATPases

Among the 39 proteins identified in the development stage I, proteins related to the secretory pathway (ER, Golgi, Golgi – ER), plasma membrane, sorting pathway (endocytosis, ESCRT), transport (vesicle-mediated transport, cytoskeleton), degradation (Autophagy-related protein 3) and to vacuolar processing as well as trafficking factors (dynamins, SNAREs, disulfide-generating enzymes and carriers, ATPase, GTPase, GTPase-activating protein and GTP binding protein) were enriched at 6 and 10 DAP (Figure 3A).

**Figure 3.**
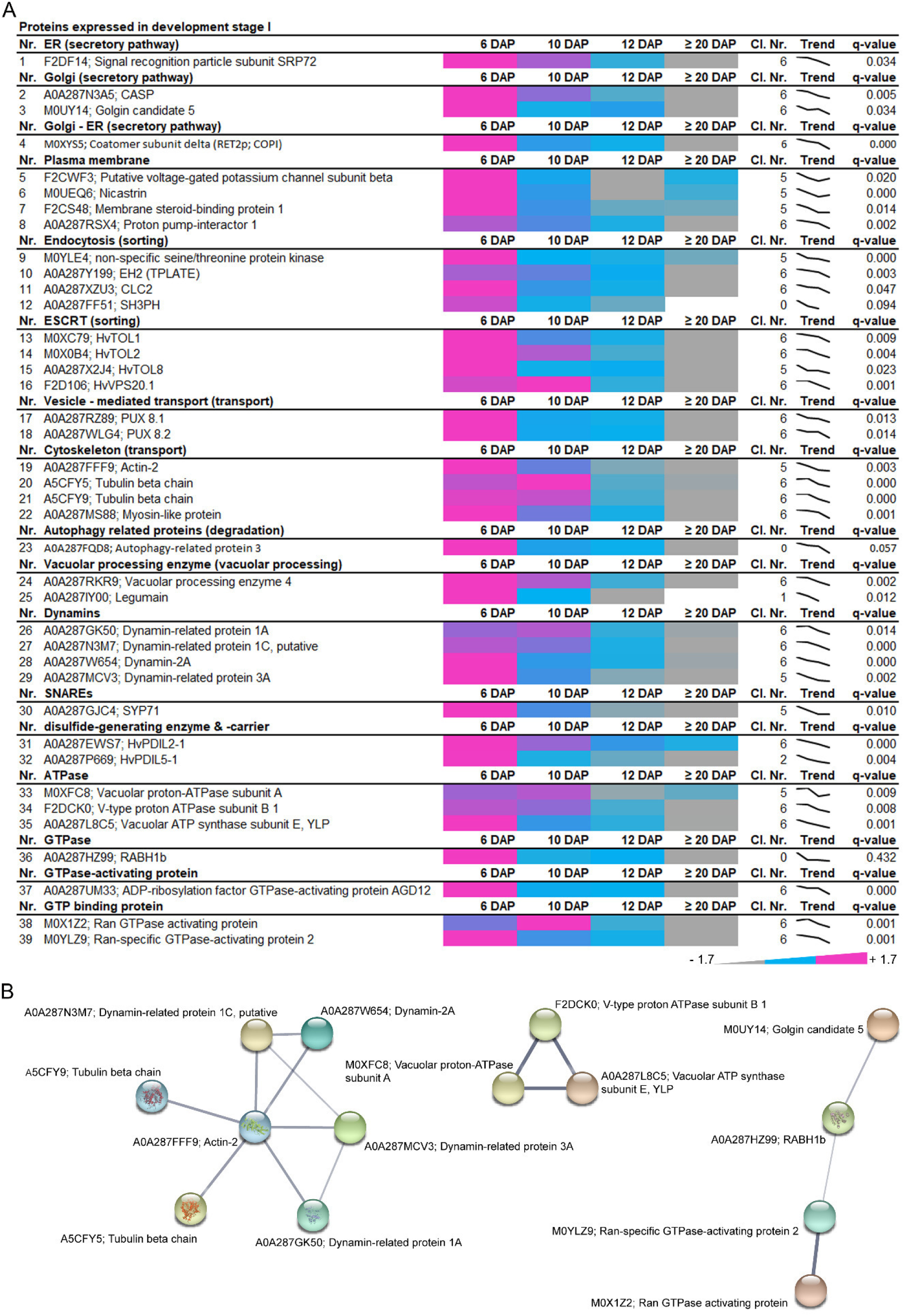
Identification of proteins that are highly abundant at development stage I of developing barley grains. (**A**) Data-matrix heat map representing z-score values of 6, 10, 12 and ≥ 20 DAP. Heat map was prepared using Microsoft Excel. Scale: grey = smallest value; blue = 50% quantile; pink = highest value. (**B**) Proteins present in this stage were analyzed using the STRING database. STRING default parameters were used (Franceschini et al., 2013), protein names are indicated.

More precisely, plasma membrane-associated proteins such as putative voltage-gated potassium channel subunit beta (F2CWF3), Membrane steroid-binding protein 1 (F2CS48), Nicastrin (M0UEQ6) and Proton pump-interactor 1 (A0A287RSX4) present decreasing abundance in concomitance to proteins associated to the endocytosis processes such as dynamins (A0A287GK50, A0A287N3M7, A0A287W654, A0A287MCV3), tubulin (A5CFY5, A5CFY9), actin (A0A287FFF9), myosin (A0A287MS88), EH2 (A0A287Y199) and CLC2 (A0A287XZU3). In line with the constant need of energy necessary for endocytosis and plasma membrane remodeling processes, three vacuolar-ATPase subunits were identified (A: M0XFC8, B1: F2DCK0, E: A0A287L8C5). All three subunits showed a decreased expression, indicating an acidification process at the early development stage (Supplemental Table 2). The identified CASP and Coatomer subunit delta (RET2p; COPI) were highly abundant at 6 DAP, pointing to an active retrograde transport as CASP is involved in tethering to Golgi cisternae of a subpopulation of COPI vesicles presumed to be involved in retrograde transport to the ER (Malsam et al., 2005; Latijnhouwers et al., 2007). Using STRING, a functional association between dynamins and cytoskeleton-related proteins was visualized (Figure 3B), both necessary for plant endocytic processes (Samaj et al., 2004; Samaj et al., 2005). Additionally, STRING revealed a functional correlation between several ATPases that were abundant at 6 and 10 DAP (Figure 3B). The functional association between Golgin candidate 5, RABH1b and further GTPases shown by STRING points to an active protein sorting in the trans-Golgi or at the TGN (Latijnhouwers et al., 2007). Taken together, the proteins that are highly abundant at 6 DAP point to the necessity of endocytosis, acidification, and sorting during the development stage I.

To investigate whether endocytosis is most active in this stage, we first performed time-lapse FM4-64 live cell imaging to track the endocytic pathway on sections prepared from 6 and 12 DAP. The endocytic tracer FM4-64 could be observed after 10 min at vesicles at the plasma membrane and intracellularly at membranes at 6 DAP, indicating endocytosis (Figure 4A). However, only the plasma membrane was labelled at 12 DAP (Figure 4A), indicating less endocytic activity at this stage. In addition to proteins involved in endocytosis, cytoskeleton-related proteins were highly abundant at 6 and 10 DAP (Figure 3). Interestingly, immunofluorescence studies of actin and tubulin with the antibodies anti-actin and anti-tubulin-α showed strong signals within PBs at 6 DAP, becoming weaker at 12 and ≥ 20 DAP, respectively (Figure 4B–D). Interestingly, a faint actin signal was observed at the plasma membrane at ≥ 20 DAP, whereas the signal of tubulin at the plasma membrane was strong at 6 DAP but was reduced at 12 and ≥ 20 DAP (Figure 4B–D). No signal could be detected in the negative controls for immunofluorescence for all stages (Supplemental Figure 5A).

**Figure 4.**
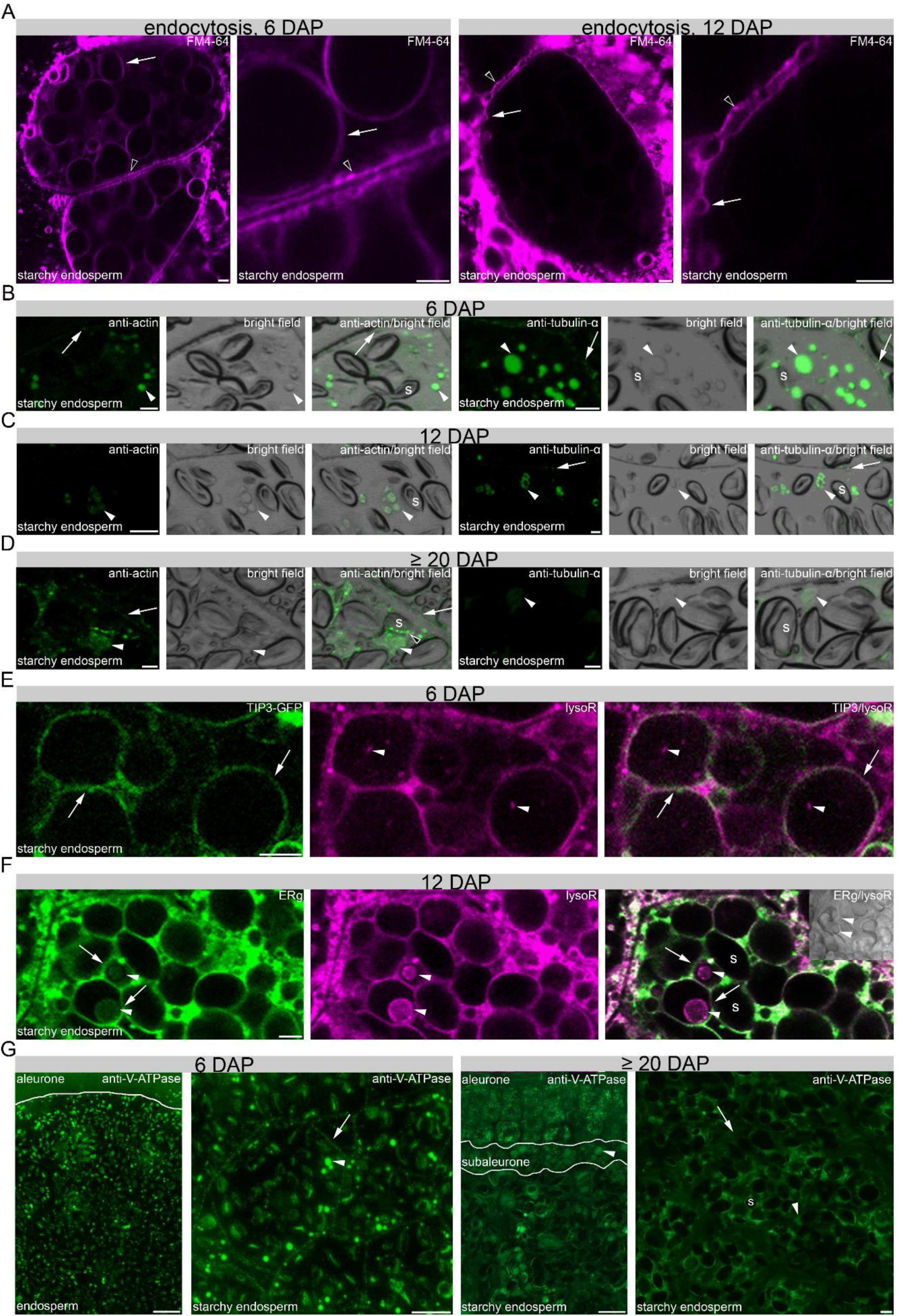
*In situ* microscopical analyses of endocytosis, cytoskeleton and acidification of PBs in development stage I. FM4-64 was internalized after 10 min of incubation, labelling intracellular membranes (arrows) and vesicles (arrowheads) at the plasma membrane at 6 DAP (**A**). At 12 DAP, FM4-64 labelled vesicles at the plasma membrane (arrowheads) and membrane structures at the plasma membrane (arrows), but not intracellular membranes. (**B–D**) Immunofluorescence studies of 1.5 µm prepared sections of 6 DAP, 12 DAP and ≥ 20 DAP using antibodies for anti-actin and anti-tubulin-α showing a strong signal at PBs (asterisks), respectively. Note the signal at the plasma membrane with anti-tubulin-α (arrow}. The fluorescence signal intensity is weaker at 12 and at ≥ 20 DAP. Note the additional signal at the periphery of the starch granule at ≥ 20 DAP using anti-actin (black-white arrowhead). (**E**) LysoTracker™ Red (lysoR) accumulation (asterisks) within TIP3-GFP labelled vacuoles (arrows) at 6 DAP. Note the lysoR-positive vesicles at the plasma membrane (arrowhead). (**F**) ER-Tracker™ Green (ERg)-labelled compartments (arrows) accumulate LysoTracker™ Red (lysoR) positive PBs (asterisks) at 12 DAP. (**G**) Immunofluorescence studies of 1.5 µm sections of 6 and ≥ 20 DAP using anti-V-ATPase antibody showing no positive signal at aleurone at 6 DAP whereas strong signal could be detected in aleurone at ≥ 20 DAP. In starchy endosperm, anti-V-ATPase antibody labels strongly PBs (asterisks) and weaker vesicles at the plasma membrane (arrowheads). At ≥ 20 DAP, the anti-V-ATPase antibody labels strongly PBs in subaleurone (asterisk), but to lesser extent in the starchy endosperm (asterisk). Note the weak labelling of vesicles at the plasma membrane (arrowhead) at ≥ 20 DAP. s = starch granule. Bars = 5 µm in A–F and 10 µm in G, except at ≥ 20 DAP where the bar represents 100 µm in the overview picture.

Recently it was shown that PBs in maize are acidified, which can be visualized by live cell imaging of fluorescent organelle markers (Ibl et al., 2018). To determine if the identified and functional associated ATPases are putatively involved in the acidification of PBs in barley, we first used LysoTracker™ Red to detect acidic compartments in the transgenic TIP3-GFP line that visualizes PSVs. At 6 DAP, acidic compartments could be observed within PSVs in the starchy endosperm (Figure 4E). Large vacuoles are most prominent in the starchy endosperm at 6 DAP (Ibl et al., 2014) when the accumulation of PBs has just started (Supplemental Figure 3) (Roustan et al., 2018), indicating that the LysoTracker™ Red-labelled compartments represent PBs. Indeed, co-labelling of ER and acidic compartments in starchy endosperm cells at 12 DAP revealed LysoTracker™ Red and ER-Tracker™ Green-positive PBs (Figure 4F).

To determine the time-dependent subcellular distribution of ATPase, we used fluorescence microscopy with anti-V-ATPase subunit epsilon antibody on sections at 6 and ≥ 20 DAP (Figure 4G). Whereas at 6 DAP a punctate structure was observed at the plasma membrane and a strong signal within PBs, a weaker labelling could be observed at the periphery of starch granules and within PBs at 12 DAP (Supplemental Figure 4B). At ≥ 20 DAP, an additional strong signal appeared in aleurone, whereas the signal was weak within PBs and at the periphery of starch granules in the starchy endosperm (Figure 4G). No signal could be detected in the negative controls for immunofluorescence for 6, 10, 12 and ≥ 20 DAP (Supplemental Figure 5B). Overall, our proteomics and *in situ* microscopic results point to a high abundance of proteins involved in endocytosis, cytoskeleton regulation and acidification of PBs at early barley grain development stage.

### Sorting-associated proteins preferentially accumulate in development stage II

Among the 15 proteins associated with development stage II (Figure 2, yellow tree), we found one peroxisome protein, PEX5 (A0A287WFD7); one cytoskeleton protein, Actin-depolymerizing factor 4 (F2DY31); proteins related to endocytosis, CHC1 (A0A287R3U8), Auxilin-related protein 1 (A0A287QP21); as well as proteins of ESCRT machinery, vacuolar processing enzymes (VPEs), GTP-binding proteins, GTPases, and ATPases (Figure 5A). Again, three ESCRT proteins, TOL3 (A0A287NWK0), SNF7.1 (A0A287R803) and SNF7.2 (A0A287XAB9) were identified, pointing to sorting processes including MVBs already detected at development stage I.

Using STRING, a functional association between the ESCRT TOL3, clathrin, and an auxilin-related protein appeared (Figure 5B). TOLs have putative clathrin-binding motifs (Korbei et al., 2013) and auxilin-related protein 1 functions as clathrin-uncoating factors (Adamowski et al., 2018), and both were described to be involved in endocytosis. Interestingly, proteins involved in sorting, transport, and degradation have been already identified in the development stage I.

**Figure 5.**
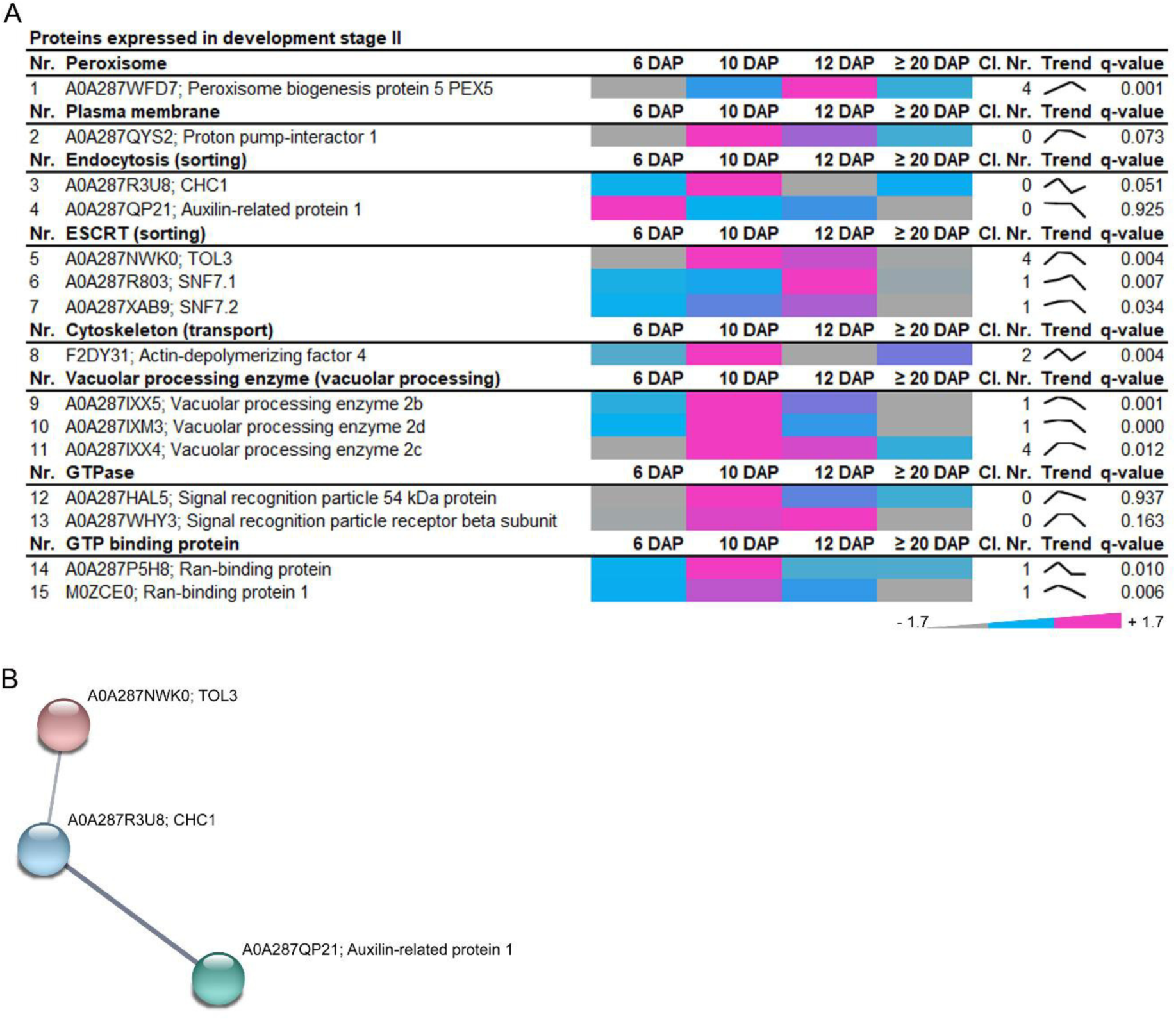
Identification of highly abundant proteins at development stage II of developing barley grains. (**A**) Data-matrix heat map representing z-score values of 6, 10, 12 and ≥ 20 DAP. Heat map was prepared using Microsoft Excel. Scale: gray = smallest value; blue = 50% quantile; pink = highest value. **(B)** Proteins present in this stage were analyzed by STRING database. STRING default parameters were used (Franceschini et al., 2013), protein names are indicated.

### Compartment-specific proteins and trafficking regulators participating in storage protein targeting, transport, and deposition are accumulating at development stage III

In total 40 proteins were associated with development stage III, most of them highly abundant at ≥ 20 DAP. Interestingly, 11 proteins involved in the secretory pathway (e.g., COPII/COPI), and 9 proteins that function in the protein sorting (e.g., VSR1, retromer, and ESCRT) were identified (Figure 6A). Additionally, proteins involved in degradation (autophagy-related 8c, vacuolar processing enzyme1) were highly abundant at ≥ 20 DAP. Among other proteins, one SNARE was identified as well as 11 trafficking regulators associated with the ATPase, GTPase or the GTP-binding group. Strikingly, endomembrane-associated proteins characterizing the stage III presented the highest interconnectivity within the STRING database compared to stages I and II (Figure 6B).

**Figure 6.**
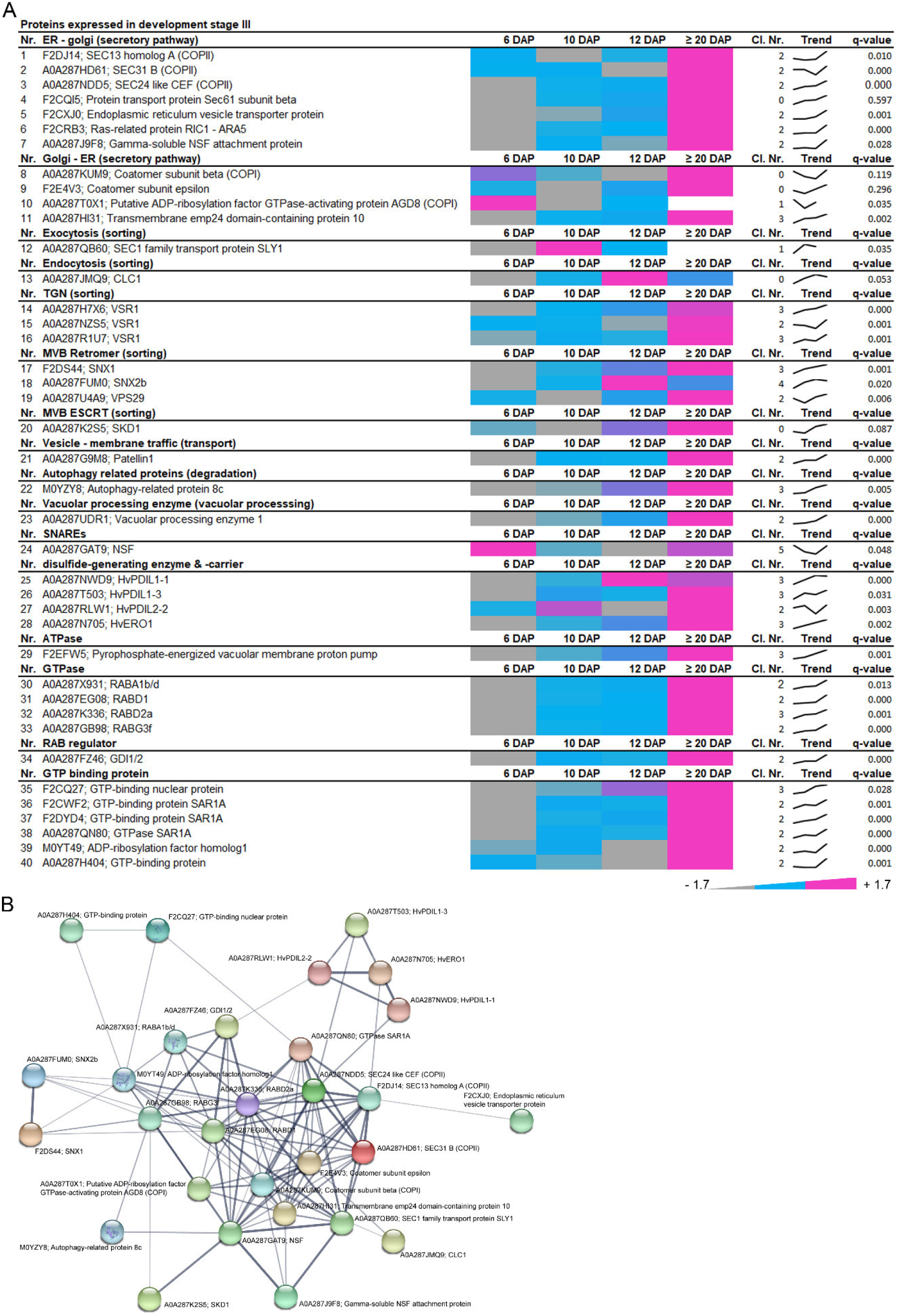
Identification of highly abundant proteins at development stage III of developing barley grains. (**A**) Data-matrix heat map representing z-score values of 6, 10, 12 and ≥ 20 DAP. Heat map was prepared using Microsoft Excel. Scale: gray = smallest value; blue = 50% quantile; pink = highest value. (**B**) Proteins present in this stage were analyzed by STRING database to analyze the gene network. STRING default parameters were used (Franceschini et al., 2013), protein names are indicated.

As expected, important ER markers such as the disulfide isomerase protein group (HvPDIL1-1 and ERO1) were found in this group, being in line with the accumulation of SSPs, described within Supplemental Figure 2. The STRING network indicates that the group of ER markers is linked with proteins associated with the secretory pathway and their associated factors such as a RAB regulator GDI1/2 (A0A287FZ46). Secretory proteins are synthesized on cytoplasmic ribosomes and finally translocated across the ER membrane through a channel formed primarily by the Sec61protein. There, chaperones support the proteins reaching their correct native conformations, which subsequently exit the ER and move to the Golgi apparatus. Figure 6B shows the central core formed by the coat protein complex (COPII) together with COPI. COPII is responsible for the ER–Golgi transport and is composed of Sar1, Sec23, Sec24, Sec13, and Sec31, while COPI regulates Golgi-ER transport and is represented here by coatomer subunits (A0A287KUM9 and F2E4V3) as well as other regulators (A0A287T0X1 and A0A287HI31). Interestingly, this core set of proteins are functional associated with diverse proteins including ESCRT related SKD1 protein (A0A287K2S5), autophagy-related protein 8c (M0YZY8), Sorting nexins, as well as regulatory factors related proteins (GTPases, GTP binding proteins).

However, VSR1, which participates in vacuolar sorting of 12S globulins and 2S albumins in Arabidopsis seeds (Zouhar et al., 2010), was strongly localized in the aleurone layer instead of the starchy endosperm at ≥ 20 DAP (Supplemental Figure 6A), indicating predominantly functional activity in the aleurone layer during development or during germination. This observation is supported by the fact that programmed cell death (PCD) of the starchy endosperm starts in development stage II around 15 DAP (Dominguez and Cejudo, 2014). Here, our electrolyte leakage (EL) assay, which indicates cell membrane integrity damage caused by PCD (Jiang et al., 2017), showed a strong increase between 12 and ≥ 20 DAP (Supplemental Figure 6B), supporting that PCD happens in the starchy endosperm at development stage II and III and subsequently indicates tissue-specific functions of proteins.

Some of the identified functional groups clustered within each development stage of barley grain development, suggesting a temporal regulation of the identified mechanisms. For example, vacuolar processing enzymes (VPEs) are known to be involved in PCD (Hara-Nishimura et al., 1998; Hatsugai et al., 2015; Radchuk et al., 2018) as well as in processing seed storage proteins in seeds (Shimada et al., 1994; Shimada et al., 2003; Wang et al., 2009). Similarly, ESCRT-related proteins contained members of each development stage.

### ESCRT-III HvSNF7 associates to MVBs at development stage I and both localize to and within PBs at development stage II and III

Proteomic analysis identified eight proteins related to ESCRT-0, ESCRT-III and SKD1 complex. Interestingly, they showed different expression patterns and subsequently belonged to different clusters (Figure 7A). Barley homologs of TOL1, TOL2, TOL8, and VPS20.1 showed a high abundance at development stage I, but a continuous decrease over the development of barley grain (Figure 7A). TOL3, SNF7.1, and SNF7.2 exhibit an expression peak during development stage II, while VPS4 continuously accumulated during the early grain development up to development stage III.

**Figure 7.**
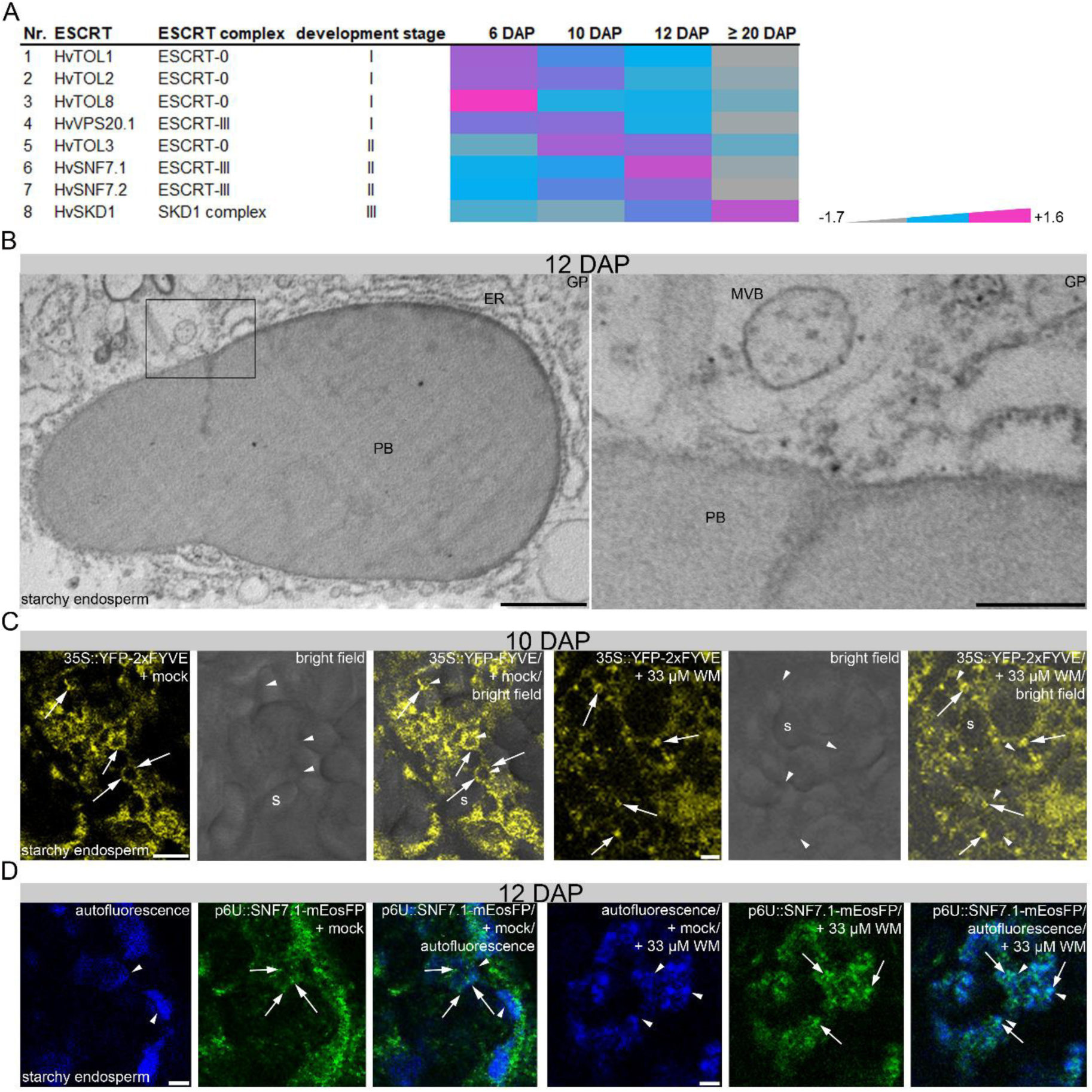
Temporally regulation of ESCRT proteins and association of SNF7.1 to MVBs associated with PBs. (**A**) Data-matrix heat map representing z-score values of 6, 10, 12 and ≥ 20 DAP. Heat map was prepared using Microsoft Excel. Scale: grey = smallest value; blue = 50% quantile; pink = highest value. **(B)** TEM section (ca. 90 nm in thickness) of starchy endosperm, ≥ 20 DAP. The zoom in on the right represents the boxed area in the overview image on the left. It displays a MVB in proximity to the PB. **(C)** Confocal live cell imaging of 35S::YFP-2xFYVE labelled MVBs (arrows) that could be detected at PBs (arrowheads) at 10 DAP. Specific punctate structures were blown up (arrows) when the sections were treated with 33 µM wortmannin (WM) for 60 minutes. (**D**) Confocal live cell imaging of p6U::SNF7.1-mEosFP labelled MVBs (arrows) that could be detected at PBs (arrowheads) at 12 DAP. Specific punctate structures were blown up (arrows) when the sections were treated with 33 µM wortmannin (WM) for 60 minutes. Autofluorescence was used to identify PBs during confocal live cell imaging as previously described in (Ibl et al., 2014). Scale = 1 µm (B), 250 nm zoom in (B), and 5 µm (C, D).

To gain insight into to expression behavior of identified ESCRT members, we analyzed the transcript levels of *TOLs, SNF7s, VPS20.1*, and *VPS4* during barley grain development by RT-qPCR (Supplemental Figure 7). RNA was isolated from whole grains harvested at 6, 10, 12 and ≥ 20 DAP, and the previously characterized most stable genes were used to normalize the *ESCRT* transcripts, as described by Shabrangy et al. (2018). A high correlation between transcript and protein abundances for all identified ESCRT members was observed, except for *VPS4*. Even though the *VPS4* transcript follows the same trend as *SNF7* transcripts and proteins, VPS4 protein increased, suggesting a delay of the response or a fine-tuning of VPS4 translation (Supplemental Figure 7). These results indicate that the expression of ESCRT is temporally regulated during barley grain development.

ESCRT originally refers to a protein–protein interaction network in yeast and metazoan cells that coordinates sorting of ubiquitinated membrane proteins into intraluminal vesicles (ILVs) of the MVB (Katzmann et al., 2001; Babst et al., 2002b; Babst et al., 2002a). So far, no observations of MVBs have been performed in barley endosperm tissues. Thus, transmission electron microscopy (TEM) analyses of 12 DAP grains were performed to study the localization of MVBs. Interestingly, an MVB was identified in proximity to a fused PB (Figure 7B).

In order to study i*n vivo* the subcellular localization of MVBs, we generated the transgenic barley line YFP-2xFYVE that functions as a fluorescent PtdIns3P-specific biosensor that can act as an endosomal marker (Samaj et al., 2004; Vermeer et al., 2006). Confocal live cell imaging revealed faint, punctate structures that were associated with PBs at 10 DAP (Figure 7C). Treatment with the phosphatidylinositol-3 (PI-3) kinase inhibitor wortmannin (WM), which targets MVBs and leads to swelling of these compartments (Tse et al., 2004; daSilva et al., 2005), indicated that the YFP-2xFYVE structures were sensitive to WM at 10 DAP (Figure 7C).

Specifically, ESCRT-III is known to be necessary for membrane remodeling that drives the biogenesis of MVBs (Hurley and Emr, 2006; Hurley and Hanson, 2010). Recent electron tomography studies in *A. thaliana* revealed that intraluminal vesicles form as large networks of interconnected or concatenated vesicles. AtSNF7 was detected in the intervesicle bridges, suggesting that ESCRT-III proteins remain trapped inside the vesicle cluster in MVBs and are finally delivered together with the cargo into the vacuole (Yoshida et al., 2013; Buono et al., 2017; Otegui, 2018). We asked whether HvSNF7 is localized at MVBs in barley endosperm. Therefore, the transgenic line SNF7.1-mEosFP driven by the hordein promotor p6U was generated to study the localization *in vivo*. Confocal live cell imaging revealed faint, small punctate signals that were associated with PBs at 12 DAP (Figure 7D). To determine the localization of HvSNF7.1 in more detail, we treated the p6U::SNF7.1-mEosFP grains with WM. Live cell imaging of 12 DAP old sections indicated swelling of SNF7.1-mEosFP positively labelled structures, indicating that SNF7.1 is localized at MVBs (Figure 7D).

We next asked if the localization of HvSNF7 is temporally regulated. To answer this question, we first studied the localization of HvSNF7 *in vivo* at the early development stage using the transgenic line p6U::SNF7.1-mEosFP. Confocal live cell imaging of the p6U::SNF7.1-mEosFP transgenic line revealed vesicular structures and few agglomerations around PBs at 6 and 10 DAP (Figure 8A). AtSNF7.1 is known to form homodimers and thus can possibly lead to agglomerations (Richardson et al., 2011). Indeed, BiFC and Y2H analyses revealed homodimerization of HvSNF7.1 (Supplemental Figures 8A, B). As live cell imaging in developing endosperm is limited to mid- and late development stages (Ibl et al., 2014), we used an anti-SNF7 antibody to analyze the localization by immunofluorescence microscopy. Immunofluorescence analyses of sections prepared from all development stages confirmed observations of small vesicular structures and few agglomerations at 6 DAP (Supplemental Figure 7C) and strong signals within PBs at 12 and ≥ 20 DAP (Figure 8B). The punctured structures that could be observed inside the PBs possibly appeared by fusing of several smaller PBs. (Figure 8B). Additionally, a weak signal could be observed at the periphery of starch granules (Figure 8B) at 12 and ≥ 20 DAP. Interestingly, faint signals of small vesicular structures could be observed at single PBs, indicating that the localization of SNF7 between 10 and 12 DAP is fluent. These findings indicate that HvSNF7 is associated with single PBs and is localized inside fused PBs.

**Figure 8.**
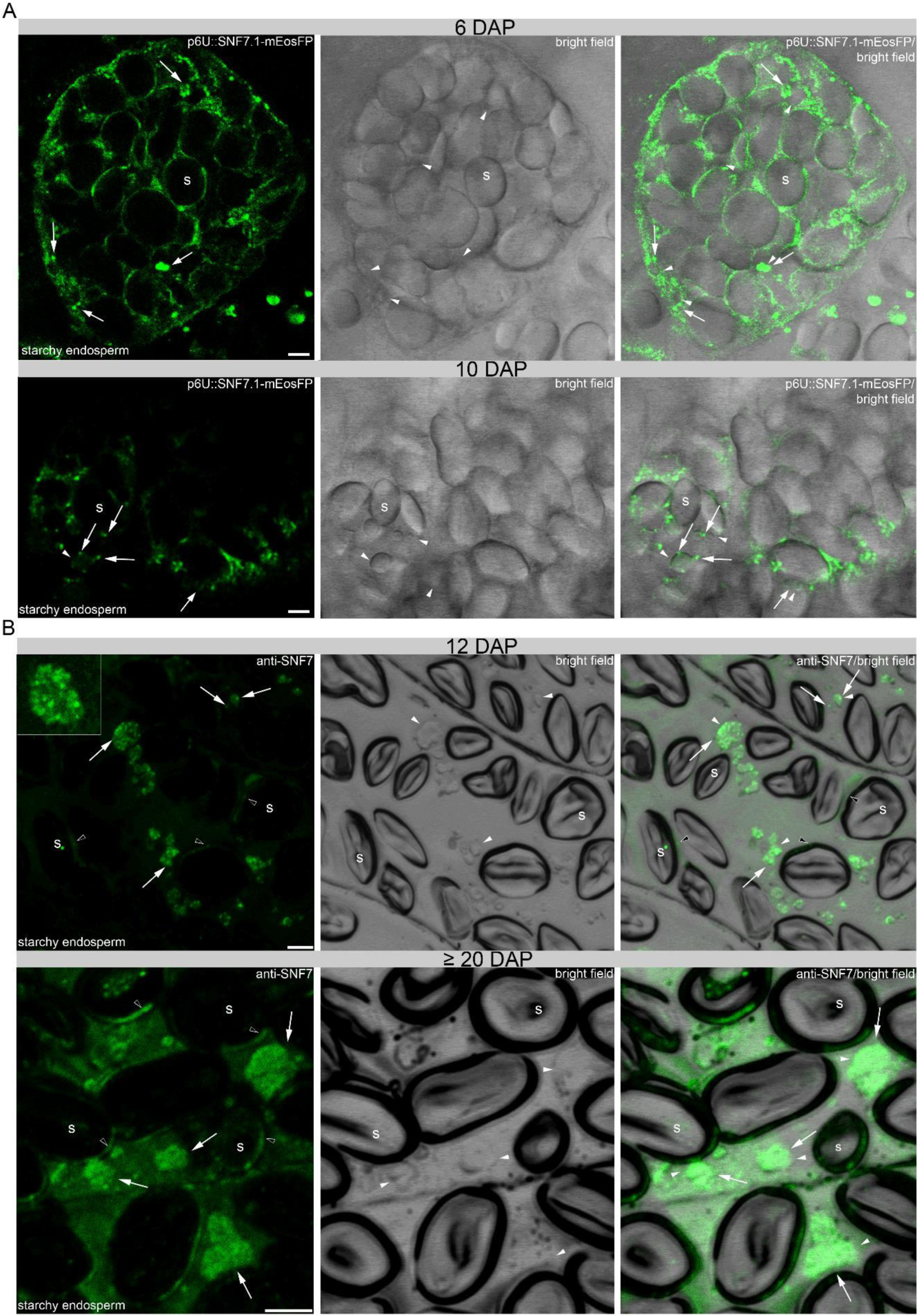
Localization of HvSNF7 in developing barley grains. (**A**) Confocal live cell imaging of p6U::SNF7.1-mEosFP at 6 and 10 DAP. Note the punctate structure (arrows) at 6 and 10 DAP around protein PBs (arrowheads). (**B**) Immunofluorescence studies of 1.5 µm sections of 12 and ≥ 20 DAP using anti-SNF7 showed weak/strong punctate structures (arrows) at PBs/within PBs (arrowheads) at 12 DAP whereas a more diffuse signal (arrows) within the PBs (arrowheads) could be detected at ≥ 20 DAP. Open arrowheads show an additional signal at the periphery of starch granules at 12 and ≥ 20 DAP. Scale = 5 µm.

## Discussion

### Proteomics and *in situ* microscopic analyses enable the mapping of the endomembrane system of developing barley endosperm

Many proteomic analyses in barley grain allow us to identify and understand the molecular composition of developing barley grain (Finnie et al., 2002; Finnie et al., 2011; Mock et al., 2018). However, less is known about proteins regulating the endomembrane system in developing barley grains. Only one of the proteins identified in our study, Membrane steroid-binding protein 1 (F2CS48), was recently characterized in barley (Witzel et al., 2018). Recent studies suggested that the endomembrane system and SSP trafficking are co-spatio-temporally regulated in developing barley endosperm between 8 and 12 DAP Recent studies have shown that the endomembrane system and SSP trafficking are spatio-temporally regulated in developing barley endosperm between 8 and 12 DAP (Ibl et al., 2014; Roustan et al., 2018). This time-dependent regulation was confirmed by PCA analysis of our proteomics data, which revealed stage-specificity of protein expression during grain development.

A detailed analysis of the 95 proteins associated with the endomembrane system associated them with three development stages: I, II and III. Altogether with previously published work, correlation of proteomics data with *in situ* microscopic analysis allowed us to provide the first temporal map of endomembrane-related proteins involved in stage-specific regulation of different endomembrane processes (Figure 9): in development stage I, prolamin and glutelin RNAs are localized to two subdomains of the cortical endoplasmic reticulum (ER), and targeted to the Golgi or to PBs by RNA-binding proteins and the cytoskeleton (Hamada et al., 2003a; Hamada et al., 2003b; Crofts et al., 2004; Crofts et al., 2010; Tian et al., 2019). Additionally, development stage I displays endocytic activities involving plasma membrane rearrangement. Such processes have been shown to be associated with specific cytoskeleton dynamics: besides the necessity of the plant cytoskeleton in vesicle trafficking and organelle movement (Thomas and Staiger, 2014), actin is required for the auxin-dependent convolution and deconvolution of the vacuole (Scheuring et al., 2016).

**Figure 9.**
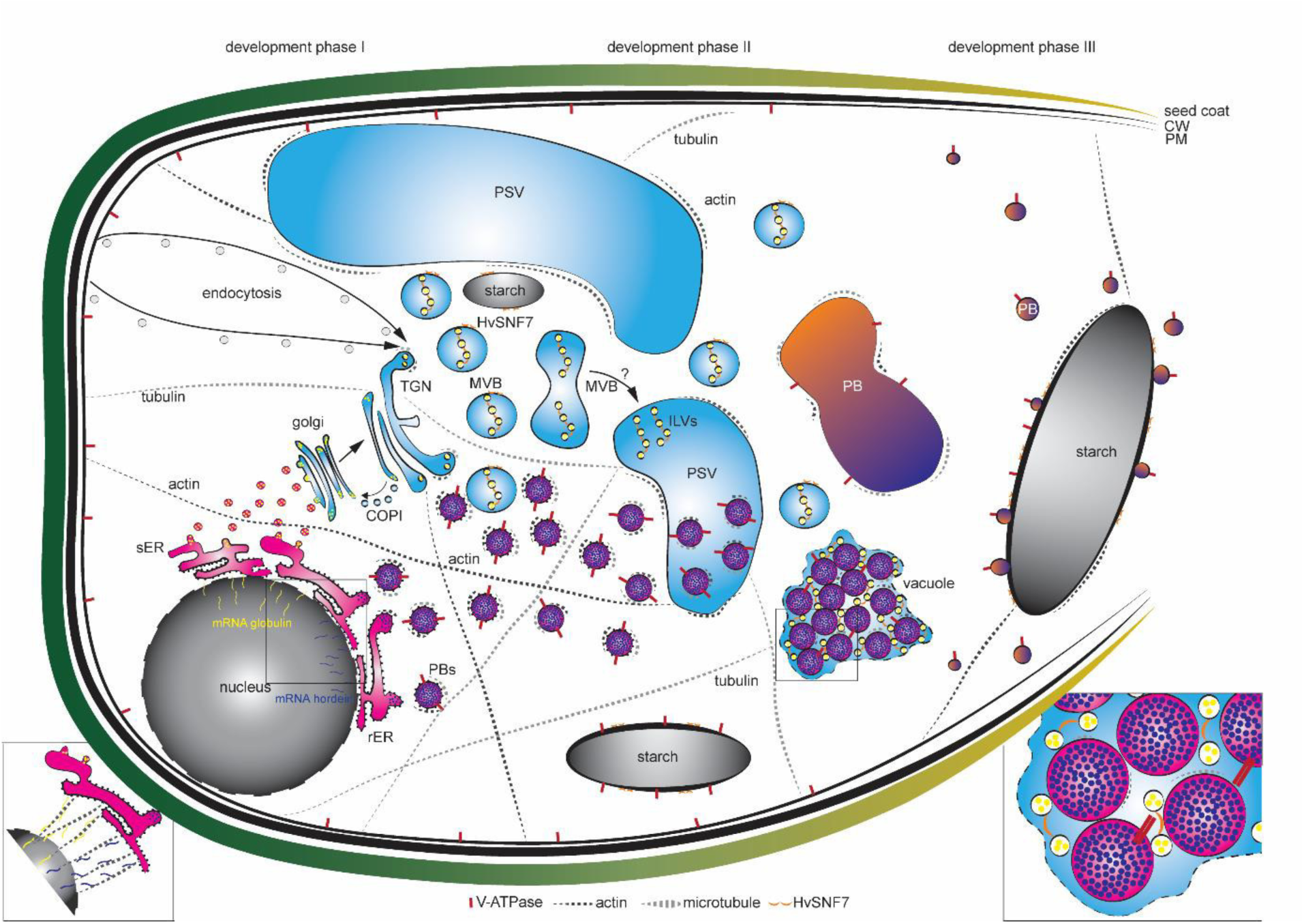
Quantitative *in situ* mapping of the endomembrane system during barley endosperm development. Quantitative proteomics and *in situ* microscopic analyses identified MVBs and HvSNF7 as putative key players for protein sorting into PBs during barley endosperm development. At development stage I, mRNA of e.g., globulin and e.g., hordein are transported by the cytoskeleton to the ER where they are entering different protein trafficking pathways (zoom in) (Hamada et al., 2003a, 2003b; Crofts et al., 2004, 2010; Tian et al., 2019). During development stage I and II, PSVs become smaller and PBs are formed, both putatively regulated by the cytoskeleton. In parallel, MVBs containing HvSNF7 appear and possibly fuse with PSVs, leading to PSVs containing HvSNF7 positive ILVs and PBs (zoom in). Between development stage II and III PSVs collapse, and PBs fuse to one big PB containing HvSNF7. At development stage III, PBs become smaller again, attaching to the protein matrix at the periphery of the starch granule. Note the additional localization of HvSNF7 at the starch granules between stage I and III. Additionally, V-ATPase localize to PBs at development stage I, acidifying PBs. V-ATPase could be further observed at starch granules during development. Schema is not in scale. PSV, protein storage vacuole; MVB, multivesicular body; ILVs, intraluminal vesicles; PB, protein body; sER, smooth ER; rER, rough ER; CW, cell wall; PM, plasma membrane

Specifically, SNAREs, actin and its associated motor protein myosin shape the vacuole by actin-dependent constrictions. While the protein accumulation starts at 6 DAP with observable PBs at 8 DAP, the size of PSVs decreases between 8 and 10 DAP. As we could observe actin as well as tubulin associated with PBs, we conclude that the cytoskeleton proteins present at early stages of barley endosperm development regulate the PBs formation/trafficking and the size of PSVs. Concurrent with the endocytosis activity, microscopic analyses indicate an increase of the acidity within PBs. During development stage II, processes associated with the sorting system seem to be highly active. Recently, MVBs were discussed to be taken up into the vacuole by autophagy (Abirached-Darmency et al., 2012). Additionally, studies in *A. thaliani* root cells have shown that central vacuoles were derived from MVB to small vacuole transition and subsequent fusions of small vacuoles (Cui et al., 2019).

Thus, we propose that MVBs loaded with HvSNF7 possibly contribute to PSV rearrangement events, resulting in large PSVs containing PBs associated with MVBs and HvSNF7, as it is known that PBs are taken up by PSVs at 10 DAP (Ibl et al., 2014). Finally, during development stage III, PBs, MVBs, and HvSNF7 were found in proximity to the protein matrix at the periphery of starch granules, while the presence of the cytoskeleton was reduced.

Here, we point out the necessity to correlate proteomics data with microscopic analysis for reasons of spatio-temporal specificity. Although the bulk of the barley grain is mainly occupied by the starchy endosperm, we cannot exclude that identified proteins can reflect different spatial activity. Thus, proteomics from dissected sections would be useful to identify tissue-specific proteins. In order to refine our analysis to a spatio-temporal level, we compared our dataset with previously published LMD-based proteomics analysis (Roustan et al., 2018). However, only four out of our 95 identified endomembrane related proteins could be detected (F2DY31, Actin-depolymerization factor 4; A0A287NWD9, HvPDIL1-1; A0A287RLW1, HvPDIL2-2; and F2CQ27, GTP-binding nuclear protein). This indicates a detection range limitation of proteomic analyses of dissected samples prepared for laser microdissection. Consequently, a combined analysis of proteomics together with microscopy appears as the most appropriate strategy.

For example, VPE4 was described to be most expressed in the pericarp of barley between 8 and 10 DAP (Radchuk et al., 2011) and to be necessary for programmed cell death execution in the developing pericarp (Radchuk et al., 2018), thereby being responsible for the grain size, starch and lipid content. As our proteomics data identified VPE4 to be most abundant at 6 and 10 DAP and subsequently to be grouped into development stage I, we assume the main function of VPE4 is in the pericarp. In contrast, VPE2b, c, and d, which are most abundant between 10 and 12 DAP and were grouped to development stage II, have been described to be involved in nucellar PCD (Radchuk et al., 2011). Additionally, besides being expressed and putatively involved in sorting processes during grain development, VSR1 was mostly expressed at ≥ 20 DAP, but mainly localized in the aleurone layer, concluding that VSR1 could be putatively involved in recycling processes during germination.

It is worth mentioning that our data provide comprehensive coverage of the endomembrane-associated proteins involved in the rearrangement of the endomembrane system and protein trafficking. However, we cannot entirely exclude additional proteins involved in these processes or organelle-specific proteins involved in other processes. For example, even though the importance of the Golgi in protein trafficking was shown previously in wheat in the transition between stage I and II (Tosi et al., 2009; Arcalis et al., 2010; Tosi et al., 2011), only two Golgi-associated proteins could be found (Golgin 5 and CASP), both grouped to development stage I.

In order to define if Golgi-associated proteins are underrepresented in our dataset, we compared our data with previously identified proteins in *A. thaliana* (Dunkley et al., 2006). Interestingly, we could explicitly identify more than 10 Golgi-resident proteins such as cell wall synthesis-associated proteins (e.g., UDP-glucuronic acid decarboxylase, A0A287H8Z0), which are parts of the glycosylation processes (Supplemental Table 2). It is possible that these identified proteins are active in aleurone, as previous data characterized the barley aleurone *N*-glycoproteome, in which numerous *N*-glycosylation sites were identified that play key roles in protein processing and secretion (Barba-Espin et al., 2014).

### How do ESCRTs and MVBs contribute to PB formation?

Although MVBs were reported to be responsible for targeting proteins to the storage vacuole in maize aleurone cells (Tian et al., 2007; Reyes et al., 2011), their existence and localization during cereal endosperm development has so far not been investigated. Here, our proteomics data revealed VPS29, SNX1, and SNX2b, proteins from the retromer complex, mediating the recycling pathway at the TGN and at the MVB (reviewed by (Paez Valencia et al., 2016). Additionally, our proteomics data identified eight ESCRT proteins (four from ESCRT-0, three from ESCRT-III and one from the SKD1 complex) quantified over all three development stages. MVBs have been suggested to arise by TGN maturation (Scheuring et al., 2011; Singh et al., 2014) with the support of Rab GTPases (Cui et al., 2016). It is worth mentioning that several Rab GTPases were detected over all three different development stages, possibly supporting MVB maturation.

Additionally, ESCRT are known to be necessary to drive the formation of ILVs in MVBs (Katzmann et al., 2001; Babst et al., 2002b; Babst et al., 2002a). Only few studies concerning ESCRT proteins are described in cereals: in maize, supernumerary aleurone layer1 (Sal1), that encodes the maize homolog of VPS46/CHMP1 (ESCRT-III associated), was found to restrict aleurone cell identity to the outer cell layer of endosperm (Shen et al., 2003). SAL1 maintains the proper plasma membrane concentration of DEFECTIVE KERNEL1 (DEK1) and CRINKLY4 (CR4), both involved in aleurone cell fate specification, by internalization and degradation of SAL1 positive endosomes (Tian et al., 2007). The rice AAA ATPase LRD6-6, which is homologous to the AAA ATPase VPS4/SKD1, was identified as an interactor with OsSNF7b/c (Os06g40620/Os12g02830) and OsVPS2.2 (Os03g43860), supporting its putative MVB-mediated vesicular trafficking function (Zhu et al., 2016). OsLRD6-6 is described to be localized at MVBs, to be required for MVBs-mediated vesicular trafficking and to inhibit the biosynthesis of antimicrobial compounds for the immune response in rice (Zhu et al., 2016).

Recently, the overexpressed ESCRT-III-associated component HvVPS60 was shown to be involved in protein targeting in developing barley endosperm (Hilscher et al., 2016). Here, seven of the identified ESCRT proteins were most expressed at development stage I and II, indicating a possible involvement in MVB body formation. The high abundances of the ESCRT-0 proteins, HvTOL1, HVTOL2 and HvTOL8 in the development stage I indicate early steps of cargo endocytic events to the vacuole: originally, nine Tom1 (target of Myb1) proteins with a domain structure similar to the VHS domain of ESCRT-0 were identified and it was speculated that these proteins are responsible to load the ESCRT machinery (Winter and Hauser, 2006). Tom1 proteins were further characterized as members of the *A. thaliana* TOL family (TOM1-LIKE), which are able to bind ubiquitin directly and participate in the endocytic trafficking of plasma membrane proteins, such as the auxin efflux facilitator PIN2 (Korbei et al., 2013). HvTOL3, which was detected at development stage II, may have an additional/different function, as it was already previously speculated that AtTOLs can participate in different pathways of distinct endosomal systems of plants (Moulinier-Anzola et al., 2014).

Our proteomics, RT-qPCR, microscopic and biochemical analyses showed a highly similar protein expression and transcription behaviour of both HvSNF7.1/2, which is reasonable as interaction studies showed heterodimerization (Richardson et al., 2011). Snf7/VPS32 induce membrane curvature at MVBs by assembling into long spiral filaments and are discussed to be involved in corralling the ESCRT cargo at the vesicle bud (Henne et al., 2012; Shen et al., 2014; Chiaruttini et al., 2015). Given the transgenic lines 35S:YFP-2xFYVE and p6U::SNF7.1-mEosFP, which showed punctate structure and is most properly labelling MVBs, we were able to detect MVBs around PBs at early and within fused PBs at later barley endosperm development stages.

Taken together, our findings suggest a putative role of MVBs and ESCRT proteins for targeting proteins to PBs. As proteins involved in protein sorting by MVBs were detected as most abundant at each specific development stage, we conclude that protein sorting by MVBs is a constant factor during endosperm development, thereby constantly delivering different cargos to PBs.

### Cargo-driven protein trafficking versus mechanically-stressed protein dynamics in barley endosperm

Grain development is marked by desiccation of the seed and the onset of dormancy. In barley, the starchy endosperm cells do not survive this desiccation and undergo PCD during grain development (Young et al., 1997), starting around 16 DAP (development stage II) (Dominguez and Cejudo, 2014). Different hallmarks such as DNA fragmentation, vacuolar tonoplast disruption and activity of caspases starting and executing apoptosis were described for starchy endosperm PCD (reviewed in (Dominguez and Cejudo, 2014; Tran et al., 2014). Besides VPEs, caspase-6-like proteolytic activity with VEIDase activity has been described to be active in autophagosomes and is involved in developmental programmed cell death in barley (Boren et al., 2006). Our proteomics results revealed two autophagy-related proteins (Atg3, Atg8c) grouped into development stage I and III, respectively. VPE1, which was described to be most expressed at late development stages in barley (Radchuk et al., 2011), grouped to development stage III. Studies in rice and in tobacco suggested that VPE1 is necessary for the maturation of rice glutelins and for PCD, respectively (Wang et al., 2009; Hatsugai et al., 2015). Additionally, crosstalk between MVB, ESCRT and autophagy was described, as the *A. thaliana* DUB, AMSH1 (Associated Molecule with the Sh3 Domain Of STAM1), and the ESCRT-III subunit VPS2.1 are important for autophagic degradation and autophagy-mediated physiological processes (Katsiarimpa et al., 2013).

Two classes of PCDs are known: vacuolar cell death and necrosis (reviewed in (van Doorn et al., 2011). Both have different morphological features such as formation of actin cables, rupture of tonoplast, accumulation of autophagosomes and small lytic vacuoles for vacuolar cell death and early rupture of the plasma membrane and shrinking of the protoplast for necrosis. Interestingly, lytic vacuoles have only been identified in the aleurone layer of barley grains (Swanson et al., 1998; Swanson et al., 2003). Given recent published data, where it was shown that the tonoplast degenerates in the starchy endosperm during development (Ibl et al., 2014) and our proteomics and microscopic data, where autophagy-related proteins, VPEs, acidification of PBs and less cytoskeleton related proteins are detected at development stage III, we identified putative key players for PCD in developing starchy endosperm already at development stage II. However, SSPs accumulate strongly at late development stages, raising the question of how proteins can be transcribed and translated during PCD. Is it possible that in the development stage II protein dynamics is driven by mechanical stress?

To conclude, our proteomic approach combined with *in situ* microscopy, which focused on endomembrane associated proteins isolated from four different stages of barley grain development, provide only a snapshot of the endomembrane system dynamics. Future studies will be needed to verify the temporal protein-protein interactions identified by the STRING analysis. Furthermore, it would be of interest to study the endomembrane system remodeling under abiotic stress conditions that are likely to affect the SSPs synthesis and/or the production of recombinant proteins. Finally, experimental proof of the involvement of cytoskeleton-related proteins, MVBs and ESCRT in sorting proteins to the PBs, ideally obtained by investigating mutant barley lines impaired in ESCRT function using proteomics and *in situ* microscopy, has yet to be provided. Nevertheless, our identified proteins that are associated with the endomembrane system are useful targets for genetic engineering to modulate SSPs accumulation and/or to improve the production of recombinant proteins.

## Abbreviations

AMSH1: associated molecule with the SH3 domain of Stam1
*A. thaliana*: *Arabidopsis thaliana*
ARF: ADP ribosylation factor
BiFC: Bimolecular Fluorescence Complementation
DAP: days after pollination
DV: dense vesicle
EC: electrical conductivity
ER: endoplasmic reticulum
ERO1: ER oxidoreductin 1
ESCRT: endosomal sorting complex required for transport
FBPA: fructose-bisphosphate aldolase
FDR: false discovery rate
GP: Golden Promise
GPC: grain protein content
HSP70: heat shock protein 70
*Hv*: *Hordeum vulgare*
mEosFP: monomeric Eos fluorescent protein
ILV: intraluminal vesicle
LC-MS/MS: liquid chromatography-mass spectrometry
MS: mass spectrometry
MVB: multivesicular body
PAC: precursor-accumulating vesicle
PB: protein body
PCA: principal-component-analysis
PCD: programmed cell death
PDI: protein disulfide isomerase
PM: plasma membrane
PSV: protein storage vacuole
RT-qPCR: real-time quantitative PCR
SAL1: supernumerary aleurone layer 1
SAM: S-adenosyl-L-methionine
SNF7: sucrose nonfermenting protein 7
SSP: seed storage protein
STRING: search tool for the retrieval of interacting genes/proteins
TEM: transmission electron microscopy
TGN: trans-Golgi network
TIP3: tonoplast intrinsic protein 3
VPE: vacuolar processing enzyme
VPS: vacuolar protein sorting
VSR1: vacuolar-sorting receptor 1
WM: wortmannin
Y2H: yeast two-hybrid

## Acknowledgement

We thank Roland Berdaguer, Doris Abraham for screening transgenic lines, cloning HvSNF7.1 for Y2H and with the TEM, respectively. EM work, including technical assistance with the TEM by Norbert Cyran, was performed at the Core Facility Cell Imaging and Ultrastructure Research, University of Vienna. We are grateful to Christiane Schwartz for RT-qPCR support and Dr. Karl Schedle from Institute of Animal Nutrition, Livestock Products, and Nutrition Physiology (TTE), University of Natural Resources and Life Sciences, Vienna for sharing the RNA lab. We thank the BOKU-VIBT Imaging-Center and Dr. Monika Debreczeny for help with LMD. We additionally thank Dr. Erika Isono for sharing the Y2H strain YT190. We thank Dr. Liwen Jiang for kindly providing the VSR1 antibody and Dr. Teun Munnik for kindly providing 2xCaMV35S::YFP-2xFYVE encoding plasmids. We thank Dr. David Teis for kindly providing the SNF7 antibody. We thank Dr. Andrea Pitzschke for kindly providing MKK4_SPYCE and MPK3_SPYNE encoding plasmids and Dr. Eszter Kapusi for kindly providing the vector p6U_phordeinD. A special thanks to the master gardeners Thomas Joch and Andreas Schröfl and their team. This work was financially supported by the Austrian Science Fund FWF (P29454-B22 and P 29303-B22) and Erasmus^+^ Job Shadowing program. We thank Dr. Alois Schweighofer and Madeleine Schnurer for critical reading of the manuscript.

---

**Supplemental Figure 1.**
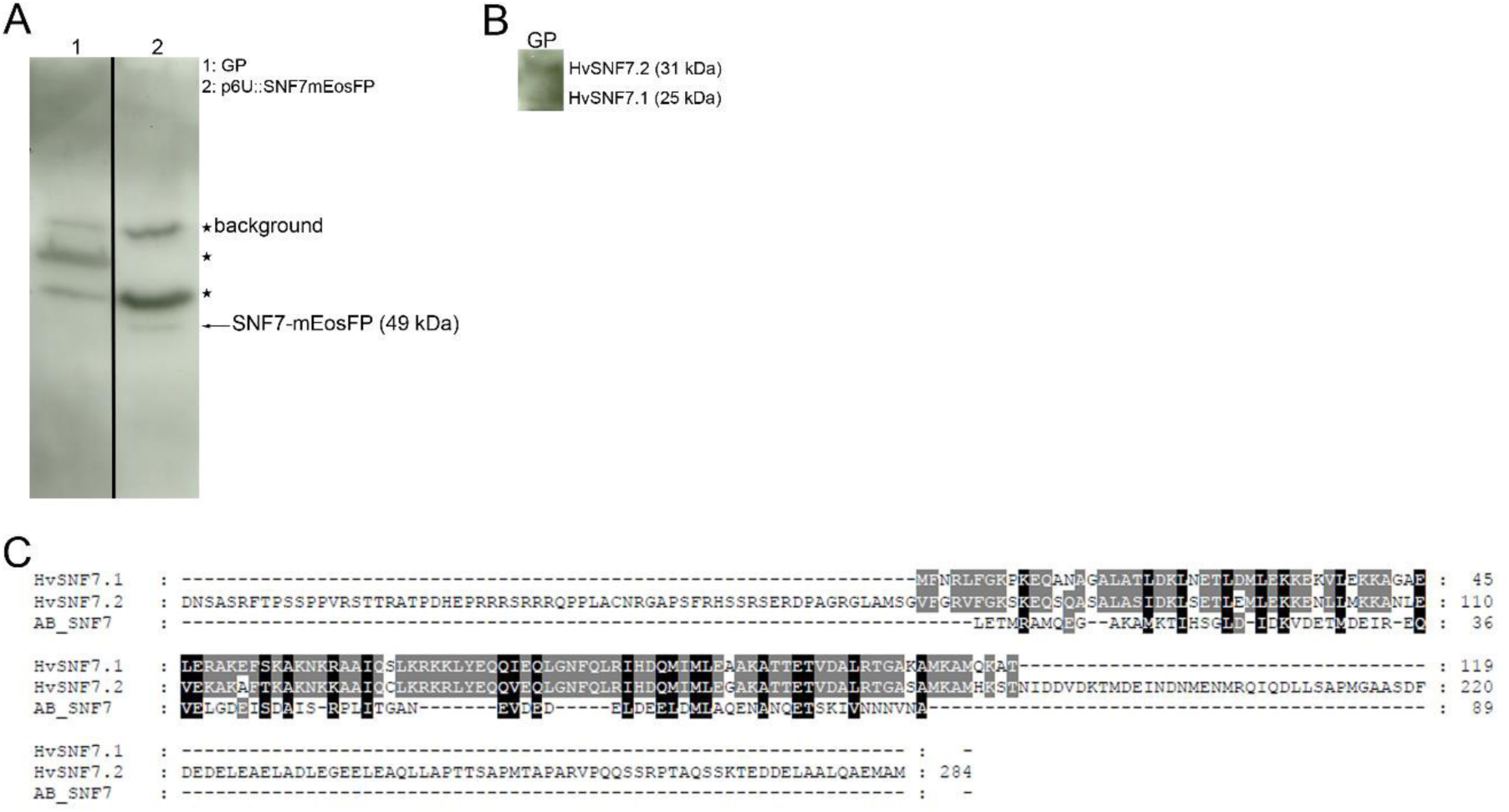
Western blot of 12 DAP barley wild type (GP) and transgenic line p6U::SNF7.1-mEosFP. (**A**) Polyclonal rabbit anti-SNF7 antibody confirm the intact fusion protein p6U::SNF7.1. (**B**) Detection of HvSNF7.1 and HvSNF7.2 using the polyclonal rabbit anti-SNF7 antibody. (**C**) Alignment of HvSNF7.1, HvSNF7.2 and the sequence of the polyclonal rabbit anti-SNF7 (Teis et al., 2010). Note the common amino acids.

**Supplemental Figure 2.**
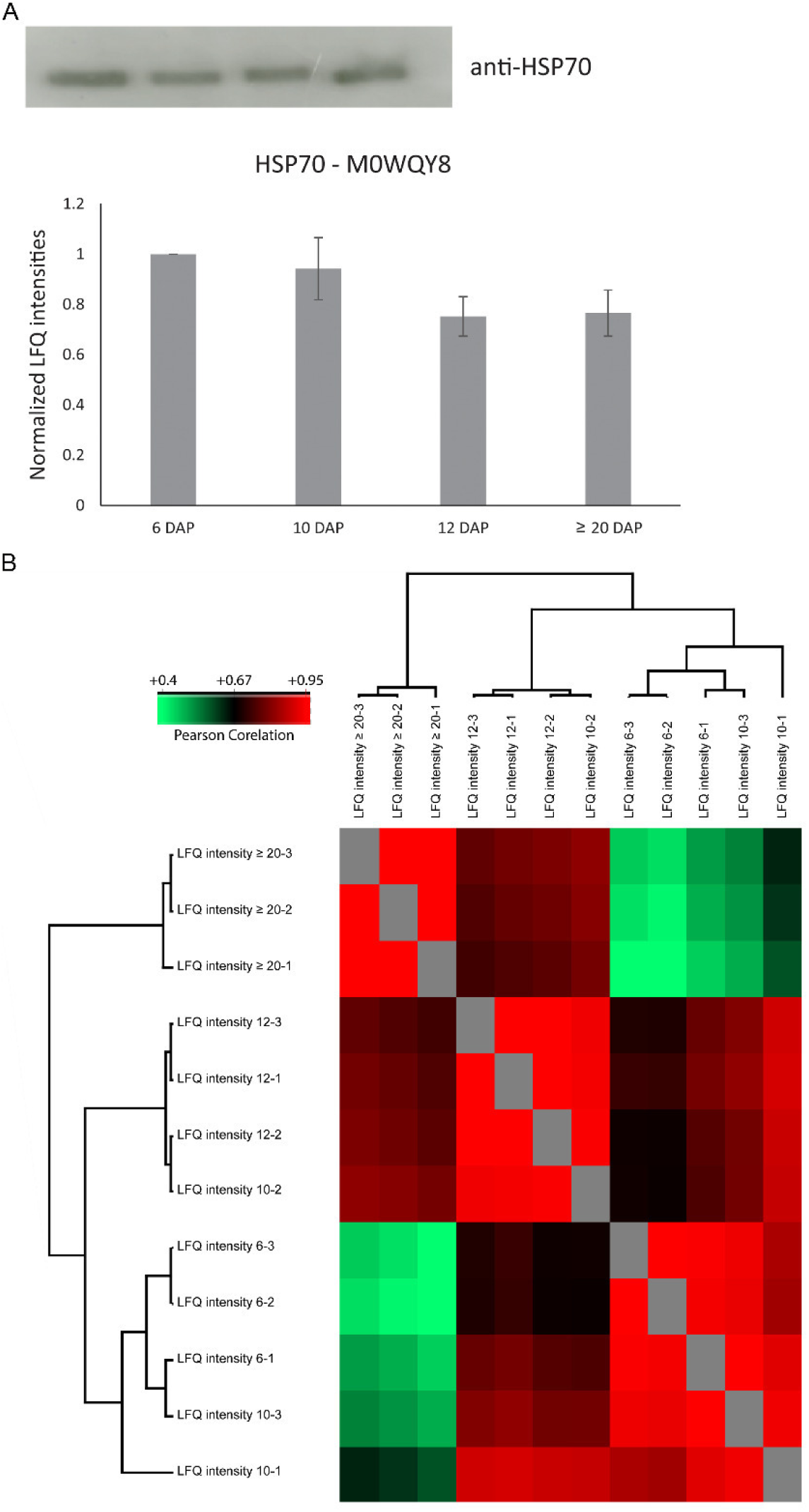
The Western blot and PCA confirm the consistency of the experimental set-up and of the obtained data. (**A**) Western blot with anti-HSP70 of protein extracts of different development stages reveals a constant signal. LFQ intensities of HSP70 (M0WQY8) at 6, 10, 12 and ≥ 20 DAP were averaged over three biological replicates. Bars represent standard deviation. (**B**) PCA shows that the measured protein abundances were highly reproducible with an average Pearson’s correlation coefficients of 0.96 between biological replicates.

**Supplemental Figure 3.**
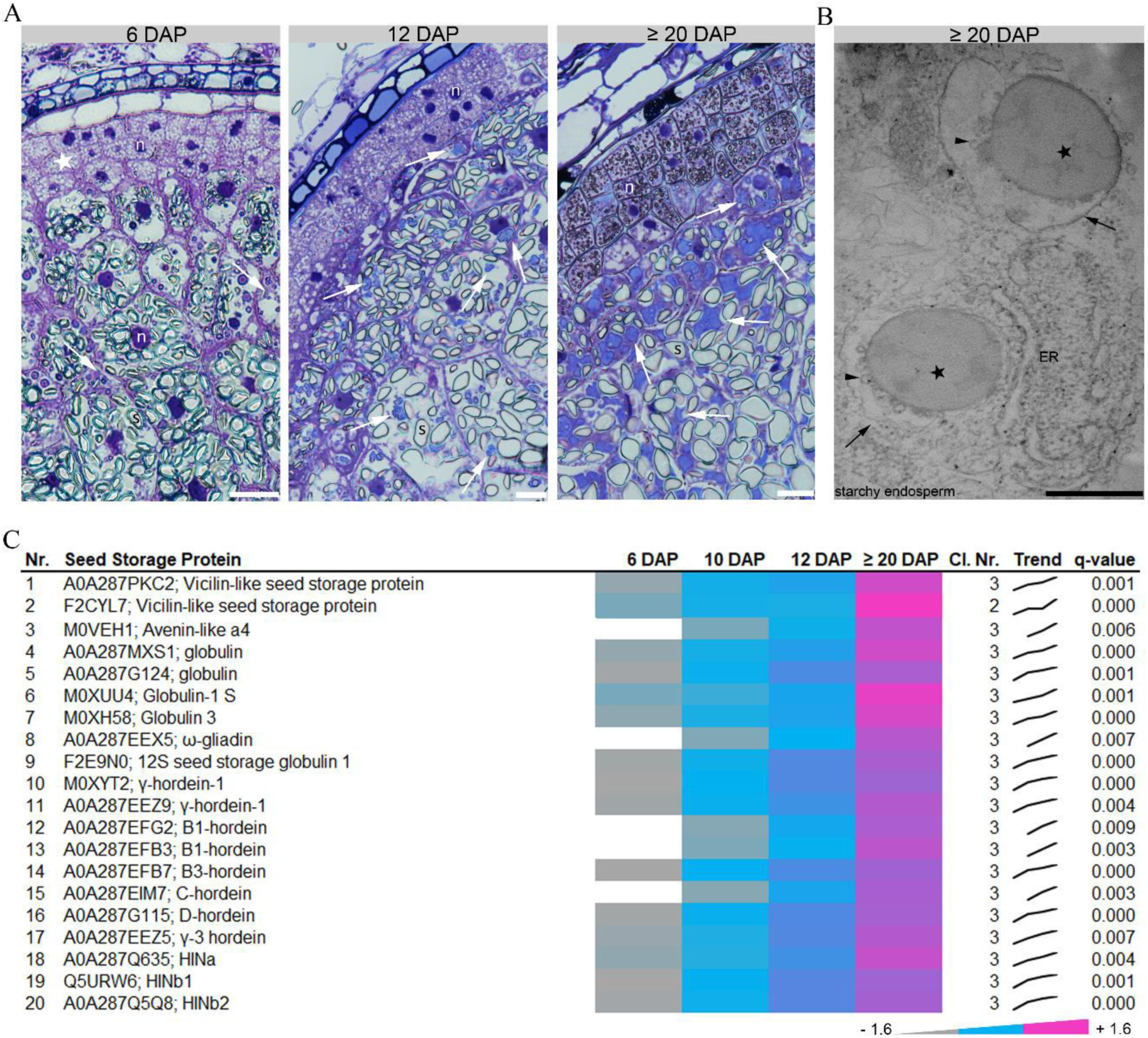
Cellularization and identification of SSPs in developing barley grains. (**A**} Toluidine blue staining of sections prepared at 6, 12 and ≥ 20 DAP. At 6 DAP, cellularization of the barley endosperm is finished and three aleurone cell layers are shown. Toluidine blue-stained compartments (arrows} were more abundant at ≥ 20 DAP compared to 12 and 6 DAP, confirming that the protein level of SSPs increase during development. Note the indicated tissues: aleurone, subaleurone and starchy endosperm. Bars = 100 µm. (**B**) Single, spherical PBs (asterisks) were observed by TEM in starchy endosperm cells engulfed by putative vacuolar membranes (arrows). Note vesicles (arrowheads) attached to the PBs. ER, endoplasmic reticulum. Bar = 0.5 µm. (**C**) Data-matrix heat map representing z-score values of 6, 10, 12 and ≥ 20 DAP. Heat map was prepared using Microsoft Excel. Scale: grey = smallest value; blue = 50% quantile; pink = highest value. Identified proteins involved in the accumulation of SSPs were highest at ≥ 20 DAP. Note that all identified SSPs could be found in cluster 3 except F2CYL7; Vicilin-like seed storage protein, that was grouped into cluster 2.

**Supplemental Figure 4.**
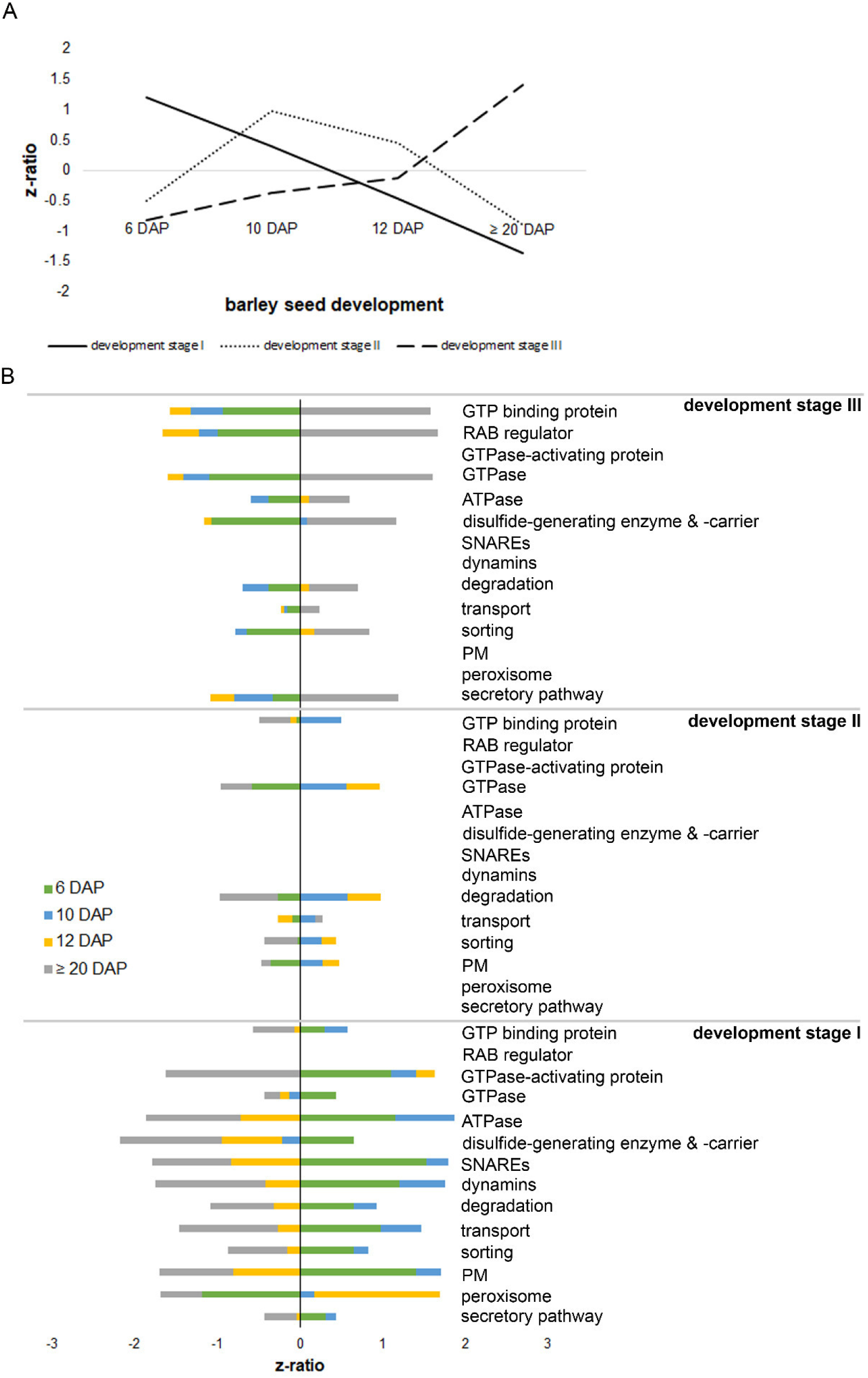
Proteins identified associated to the endomembrane system are clustering into three different development stages. (**A**) Mean of z-score values of all proteins identified in the different development stages visualizing that development stage I, II, and III behave differently during endosperm development. (**B**) Mean of z-score values of spatio-functional active proteins within development stage I, II and III.

**Supplemental Figure 5.**
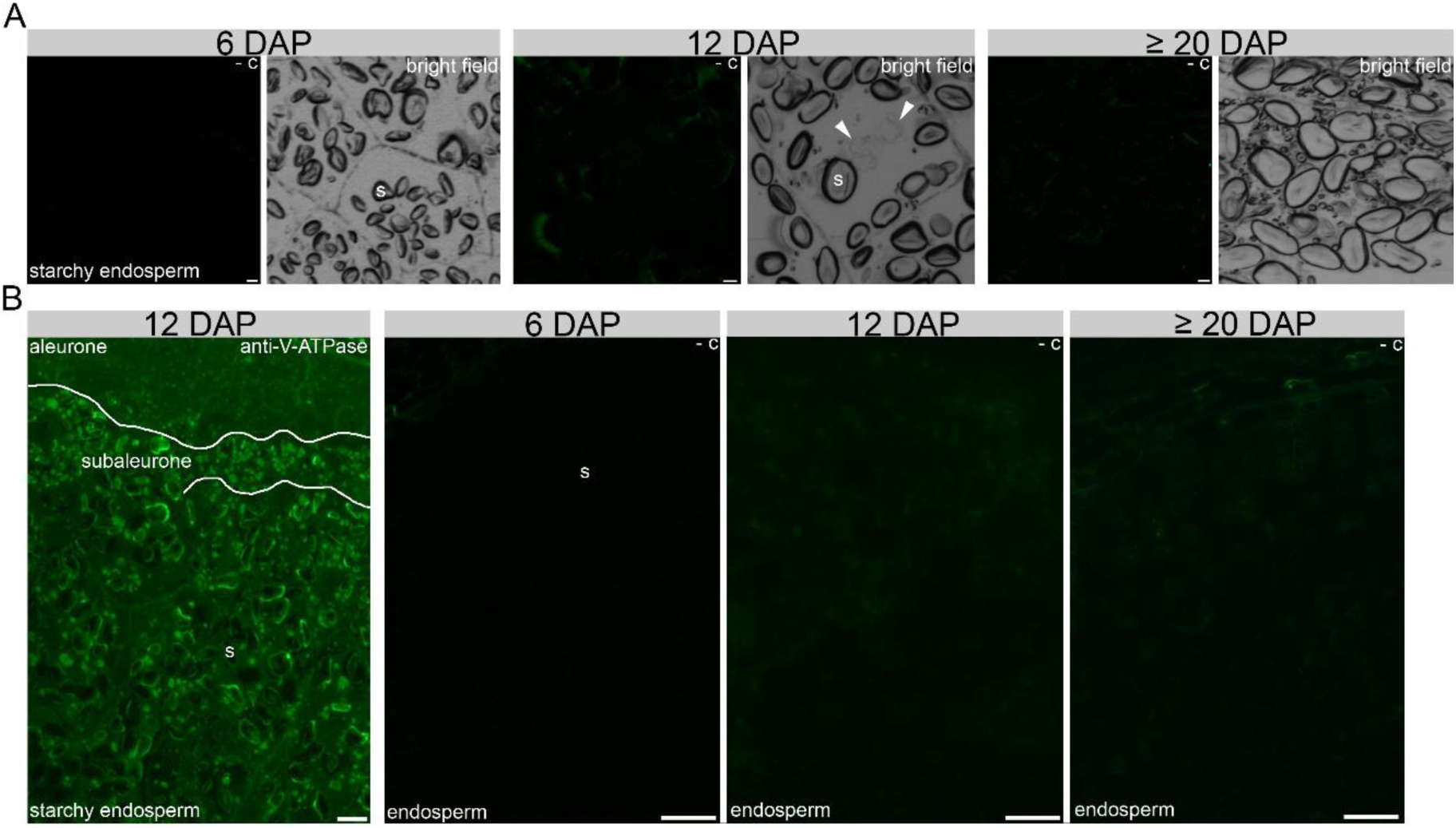
Immunofluorescence of sections prepared from 6, 12 and ≥ 20 DAP. (**A**) Negative controls from the anti-actin and anti-tubulin-α immunofluorescence assay from section prepared of 6, 12 and ≥ 20 DAP analyzed by confocal microscopy. (**B**) Immunofluorescence of anti-V-ATPase from section prepared of 12 DAP and the corresponding negative controls from section prepared of 6, 12 and ≥ 20 DAP analyses by fluorescence microscopy. Bar = 100 µm.

**Supplemental Figure 6.**
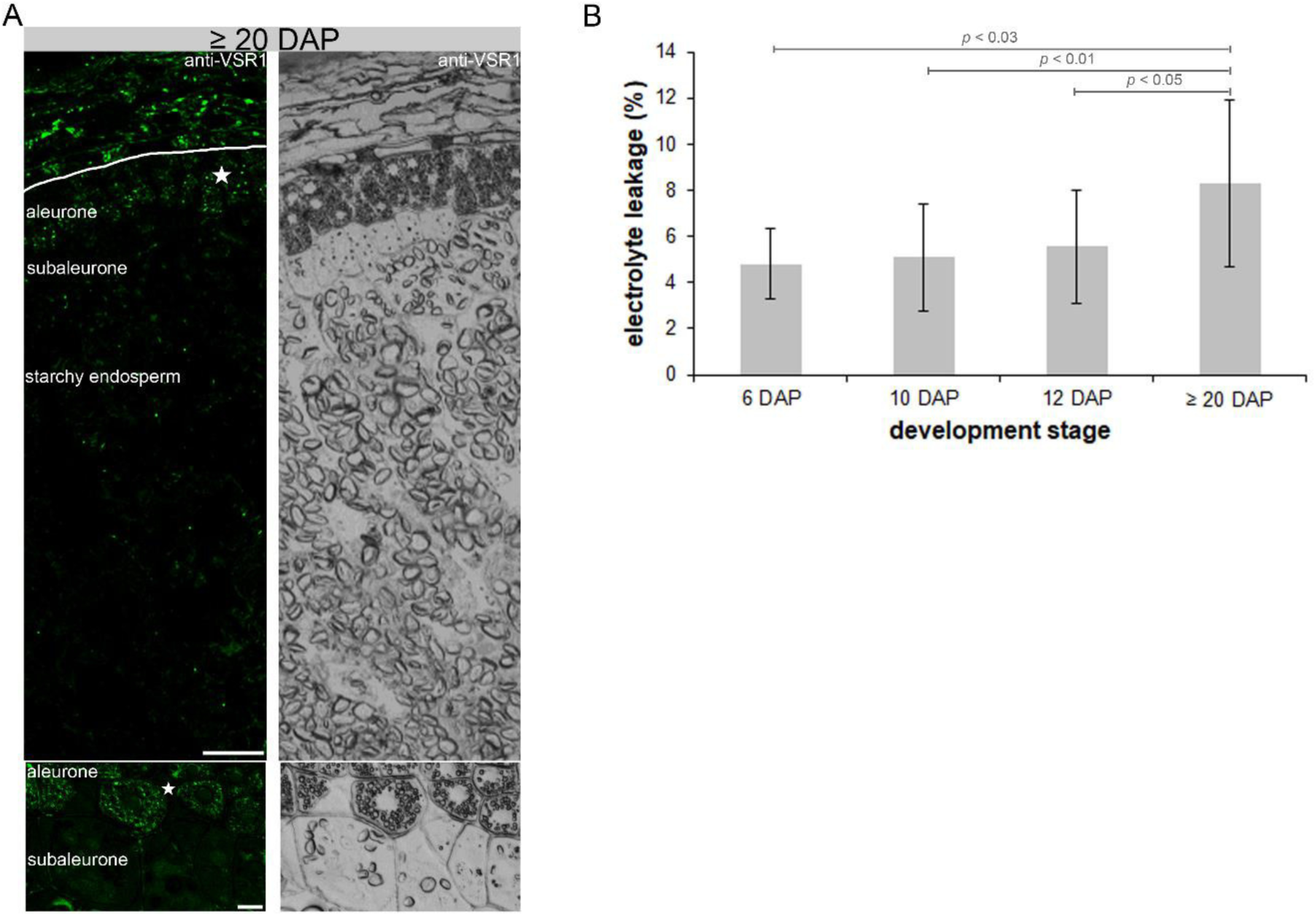
VSR1 localization and EL assay. Sections of ≥ 20 DAP were incubated with anti-VSR1 antibody and analyzed by confocal microscopy (**A**). Asterisks show the strong signal in the aleurone layer. Note the zoom at the bottom. Bar = 50 µm and 10 µm, respectively. (**B**) EL assay including different development stages. Data represent means of three biological replicates. For statistical analyses we performed a Student’s t-test (n = 3). Bars represent standard deviation. *p-*values are indicated.

**Supplemental Figure 7.**
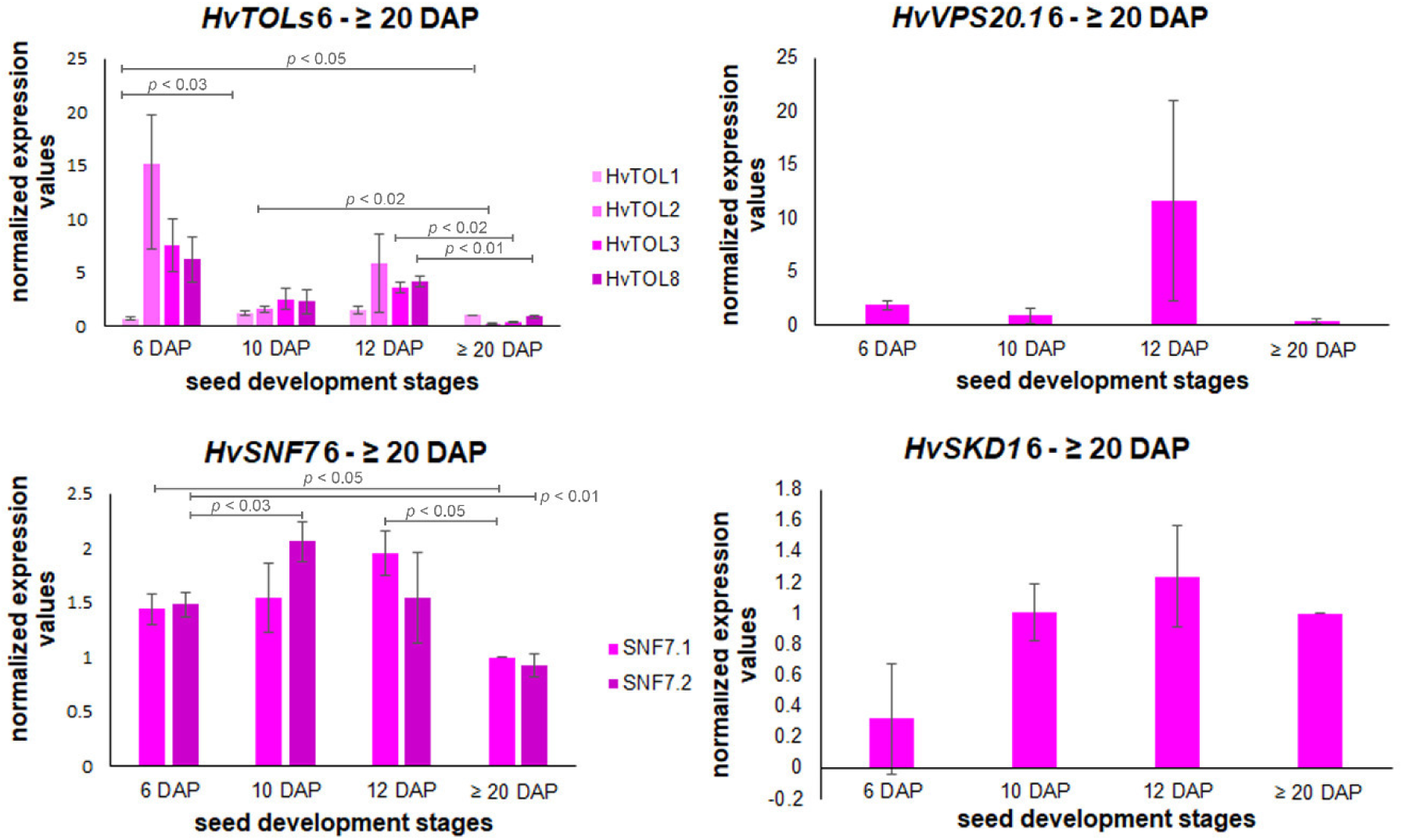
Temporal quantification of *HvESCRT* transcripts in developing barley grain. Bar graphs describe the average over three biological replicates of the normalized transcripts from *HvTOL1/HvTOL2/HvTOL3/HvTOL8, HvVPS20.1, HvSNF7.1*/*HvSNF7.2* and *HvSKD1* at 6, 10, 12 and ≥ 20 DAP. For statistical analyses we performed a Student’s t-test (n = 3). Bars represent standard deviation; *p-*values are indicated.

**Supplemental Figure 8.**
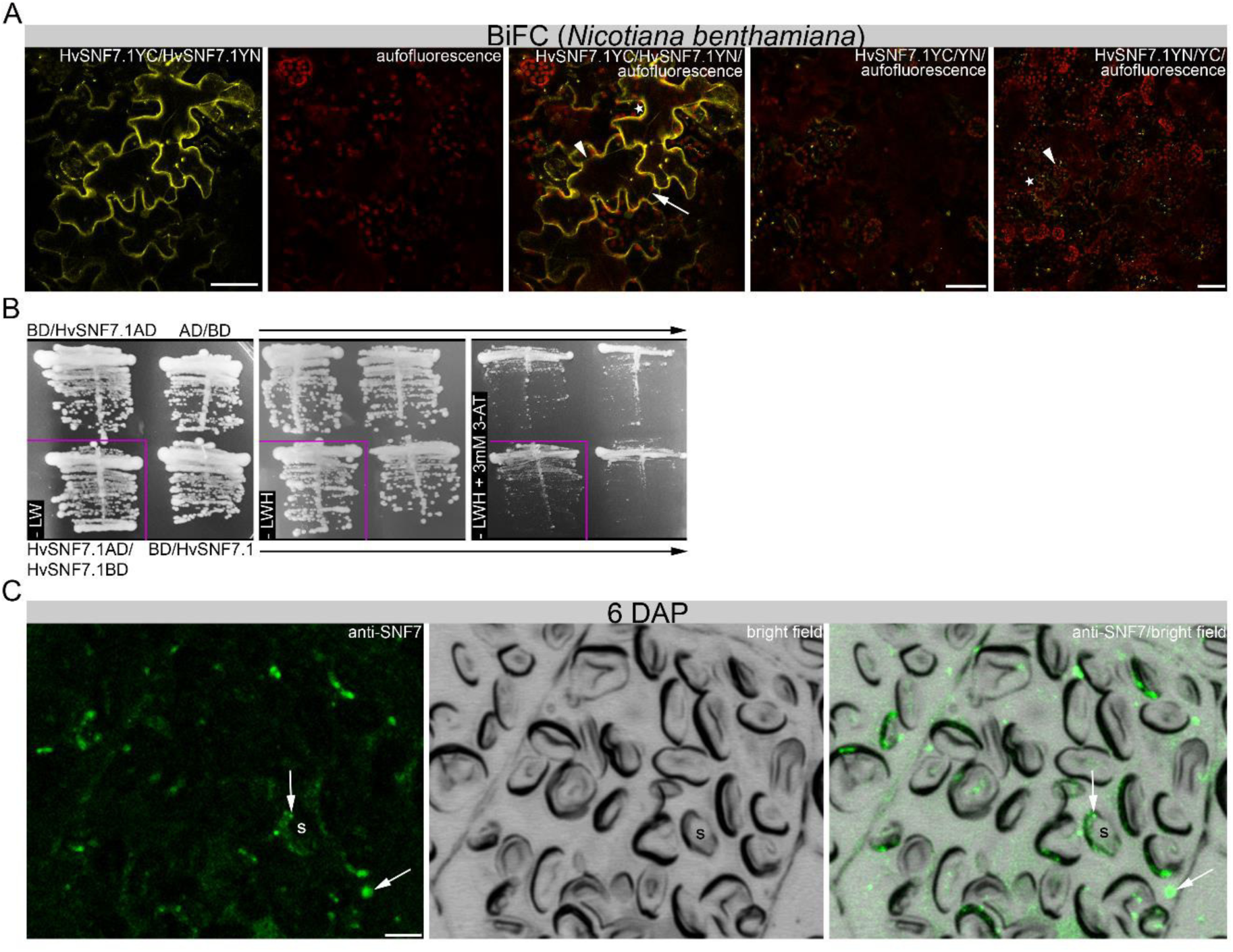
Homodimerization of SNF7.1 and immunofluorescence of SNF7. Homodimerization of SNF7.1 shown by BiFC (**A**) and Y2H (**B**). (**C**) Immunofluorescence of SNF7 at 6 DAP. Note positive labelling of vesicles (arrows) at the periphery of starch granules and associated with PBs (arrowheads). Bar = 5 µm.

